# The role of traction in membrane curvature generation

**DOI:** 10.1101/157313

**Authors:** H. Alimohamadi, R. Vasan, J.E. Hassinger, J.C. Stachowiak, P. Rangamani

## Abstract

Curvature of biological membranes can be generated by a variety of molecular mechanisms including protein scaffolding, compositional heterogeneity, and cytoskeletal forces. These mechanisms have the net effect of generating tractions (force per unit length) on the bilayer that are translated into distinct shapes of the membrane. Here, we demonstrate how the local shape of the membrane can be used to infer the traction acting locally on the membrane. We show that buds and tubes, two common membrane deformations studied in trafficking processes, have different traction distributions along the membrane and that these tractions are specific to the molecular mechanism used to generate these shapes. Furthermore, we show that the magnitude of an axial force applied to the membrane as well as that of an effective line tension can be calculated from these tractions. Finally, we consider the sensitivity of these quantities with respect to uncertainties in material properties and follow with a discussion on sources of uncertainty in membrane shape.

## Introduction

Cell shape plays an important role in regulating a diverse set of biological functions including development, differentiation, motility, and signal transduction (McMahon and Gallop, 2005; Roux et al., 2005; Neves et al., 2008; Rangamani et al., 2013; Aimon et al., 2014). Additionally, the ability of cellular membranes to bend and curve is critical for a variety of cellular functions such as membrane trafficking processes, cytokinetic abscission, and filopodial extension (Mukherjee and Maxfield, 2000; Mattila and Lappalainen, 2008). In order to carry out these functions, cells harness diverse mechanisms of curvature generation like compositional heterogeneity (Baumgart et al., 2003; Römer et al., 2007), protein scaffolding (Karotki et al., 2011a; Kirchhausen, 2012), insertion of amphipathic helices into the bilayer (Ford et al., 2002; Lee et al., 2005), and forces exerted by the cytoskeleton (Giardini et al., 2003; Carlsson, 2018) (Fig. 1). Reconstituted and synthetic membrane systems also exhibit a wide range of shapes in response to different curvature-inducing mechanisms as seen from steric pressure due to protein crowding (Lipowsky, 1995; Stachowiak et al., 2012; Derganc and Čopič, 2016).

**Figure 1:**
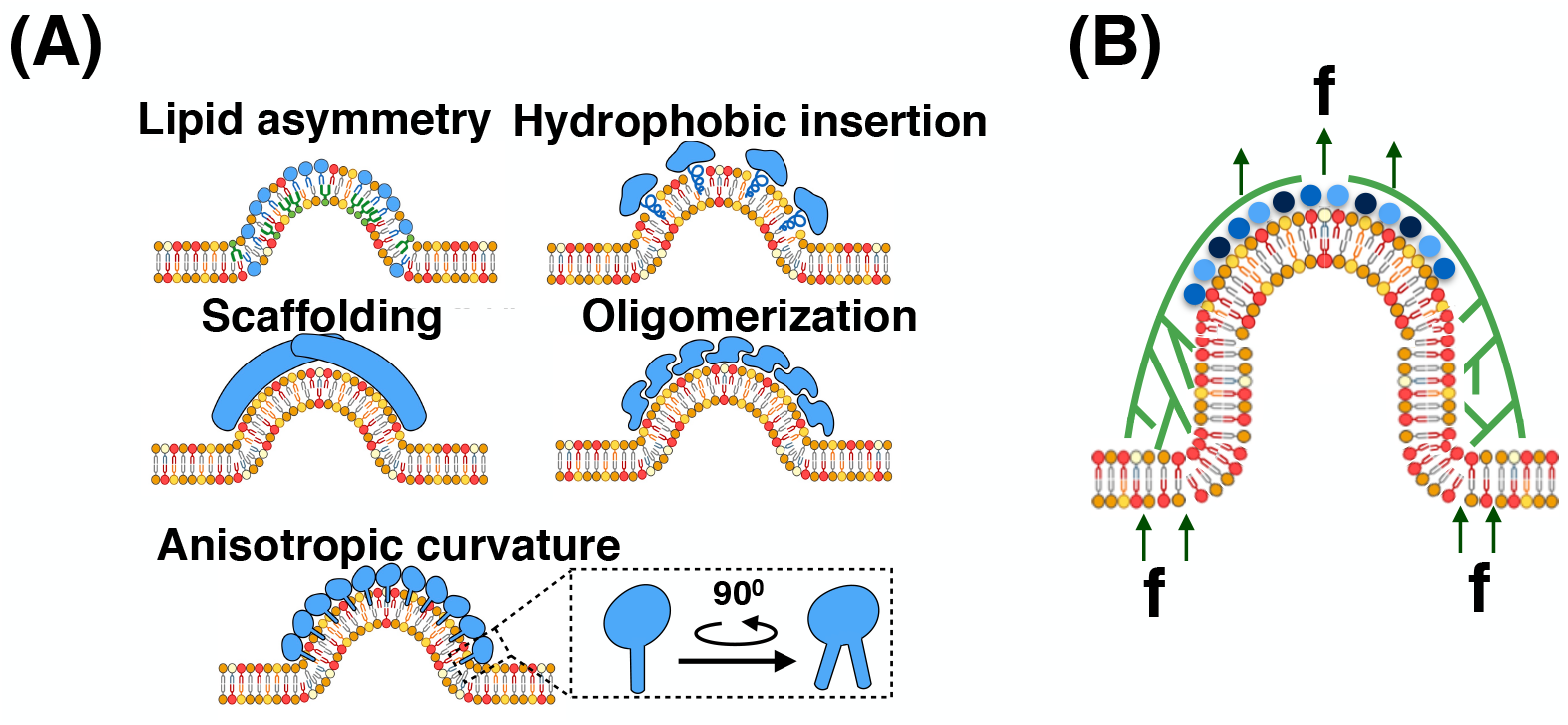
Curvature generation in biological membranes (Chabanon et al., 2017). Membrane curvature is controlled by different physical inputs including (A) protein-induced spontaneous curvature and (B) forces exerted by the cytoskeleton.

It is well-known that these various mechanisms of curvature generation induce surface stresses; expressions for these stresses have been derived using either variational methods (Jenkins, 1977; Capovilla and Guven, 2002b, 2004) or by using auxiliary variables that enforce geometric constraints (Guven, 2004; Fournier, 2007). These studies have established the physics underlying membrane stresses and clearly explained how these traction forces can be interpreted in linear deformations and in idealized geometries (Guven, 2004; Fournier, 2007). However, many physiologically relevant membrane shapes display large curvatures (Farsad and De Camilli, 2003; Kozlov et al., 2014), non-linear deformations (Holzapfel et al., 1996; Einstein et al., 2003), and heterogeneous membrane composition (Lingwood and Simons, 2010; Busch et al., 2015). How stresses are distributed along such shapes is not yet fully understood. In this article, we discuss how theory can help us evaluate membrane stresses based on the observed shape.

### Shape as a reporter of force

Many biomechanics textbooks present the postulate that the relationship between the applied load and the resulting deformation can be obtained if a constitutive relationship between the stress and strain of a material is given (Mofrad and Kamm, 2010; Phillips et al., 2012; Fung, 2013). Indeed, the idea that shape can be considered a reporter of the applied force is an idea as old as continuum mechanics (Todhunter, 1886). A classical example illustrating how shape can be used as a reporter of force in biology can be understood by studying the shape of a vesicle or a cell using micropipette aspiration (Hochmuth, 2000; Lee and Liu, 2014). This method is used to calculate the tension of bilayer membranes in vesicles and cortical tension in cells through Laplace’s law. Since the pressure applied by the micropipette is known, tension can be calculated using a force balance at the membrane.

Lee *et al.* suggested that membrane shape itself acts as a reporter of applied forces (Lee et al., 2008) and calculated the axial force required to form membrane tethers in optical tweezer experiments based on shape, given the material properties of the membranes (See Fig. 2 in (Lee et al., 2008)). They showed that the calculated value of force was in excellent agreement with their experimental measurements. Separately, Baumgart and colleagues showed that the Gaussian modulus has a strong effect on membrane budding in phase-separated vesicles and its magnitude can be obtained by analyzing the geometry of the vesicle (Baumgart et al., 2005).

**Figure 2:**
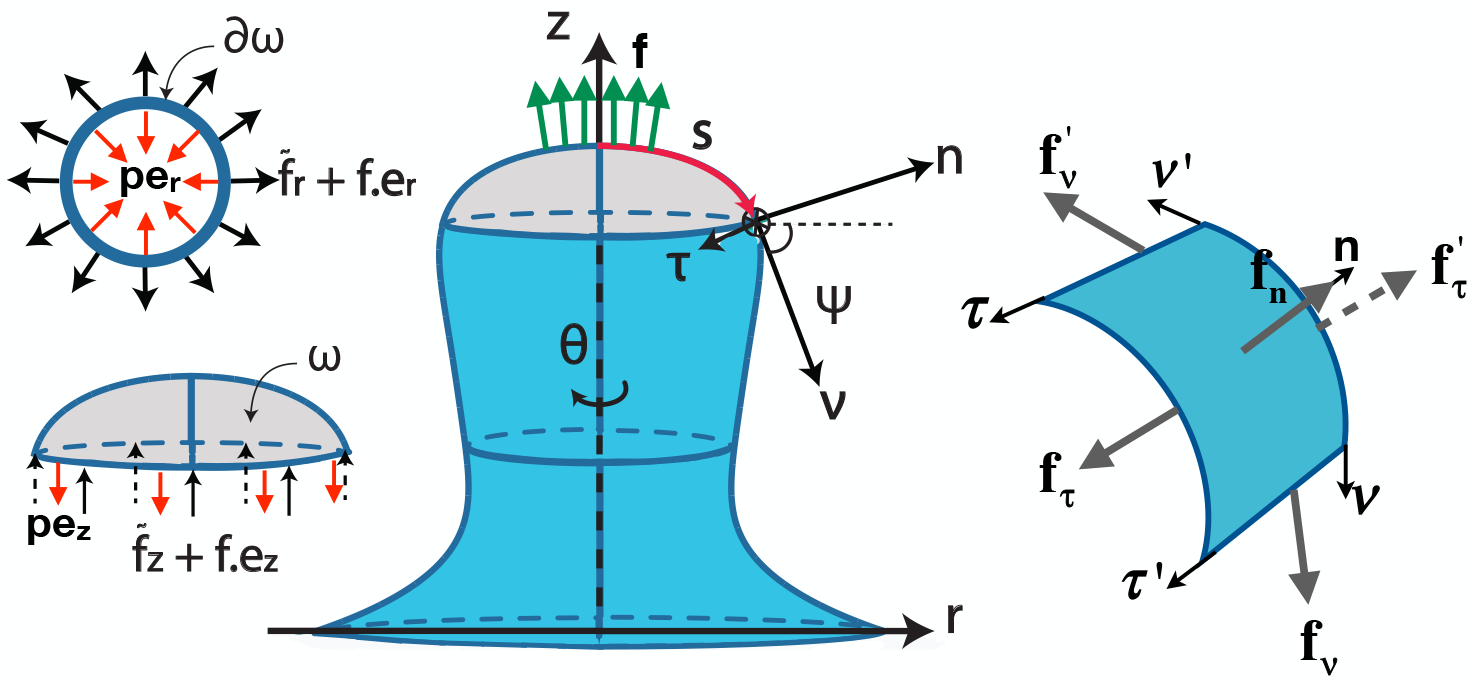
A schematic representing the axisymmetric coordinate system used for calculating curvature and traction. Inset shows that pressure opposes traction and external force in both the radial and axial directions.

An additional layer of complexity in how shape and forces are related arises through the heterogeneous composition of the lipid bilayer in cells. Most protein binding to cellular membranes are local processes (Kishimoto et al., 2011; Karotki et al., 2011b; Buser and Drubin, 2013). Even in *in vitro* studies, several groups have shown that protein adsorption on lipid domains can alter the lateral pressure profile on the bilayer and induce tubulation (Stachowiak et al., 2012; Lipowsky, 2013; Zhao et al., 2013). Recently, theoretical studies have shown that adsorbed proteins give rise to spontaneous surface tension (Lipowsky, 2013; Rangamani et al., 2014b). Therefore, there is a need to understand how applied forces and membrane heterogeneity can regulate the local stresses on the membrane. Going beyond the approximation of tension using Laplace’s law, we sought to understand the local stresses in tubes and buds – two geometries that are critical to many cellular phenomena. Using the well-established Helfrich model (Helfrich, 1973; Bassereau et al., 2014) for membrane bending as a framework, we illustrate how local forces can be understood from the shape of the membrane. We close with an extended discussion of how advances in image analysis and measurement of material properties can aid in our understanding of how traction can be calculated from the curvature of the membrane.

## Local stresses in the membrane: governing equations

### Surface stress tensor and traction calculation

A general force balance for a surface *ω*, bounded by a curve *∂ω*, is (Fig. 2)

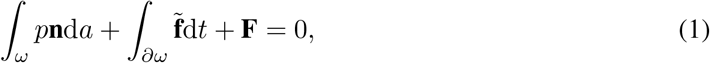

where *t* = *r*(*s*)*θ* is the length along the curve of revolution perimeter (see Fig. 2), *p* is the pressure difference across the membrane, 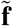 is the traction along the curve of revolution *t* and **F** is any externally applied force on the membrane. Along any circumferential curve on the membrane at constant z, the traction is given by (Agrawal and Steigmann, 2009a)

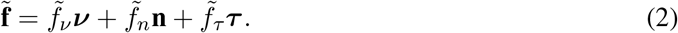

The values of *f_v_*, *f_n_* and *f_τ_* will depend on the particular form of strain energy we choose to depict the membrane properties (See Fig. 2 for definitions of the forces and the vectors). We choose the Helfrich Hamiltonian as the constitutive relationship in this case and use a modified version that includes spatially-varying spontaneous curvature *C*(*θ^α^*), (Steigmann, 1999; Agrawal and Steigmann, 2009a; Hassinger et al., 2017),

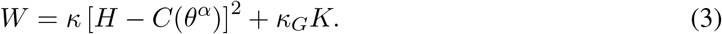

where *W* is the energy per unit area, *κ* is the bending modulus, *H* is the local mean curvature, *κ_G_* is the Gaussian modulus, *K* is the local Gaussian curvature and *θ^α^* denotes the surface coordinates. This form of the energy density accommodates the local heterogeneity in the spontaneous curvature *C*. Note that *W* differs from the standard Helfrich energy by a factor of 2, which is accounted for by using the value of *κ* to be twice that of the standard bending modulus typically encountered in the literature (See Table S1 for notation). A more in-depth investigation of the role of anisotropic spontaneous curvature using a version of the Helfrich energy that includes deviatoric curvature can be found in the Supplement (Eq. S11, (Iglic et al., 2006; Lokar et al., 2012)).

**Table 1:**
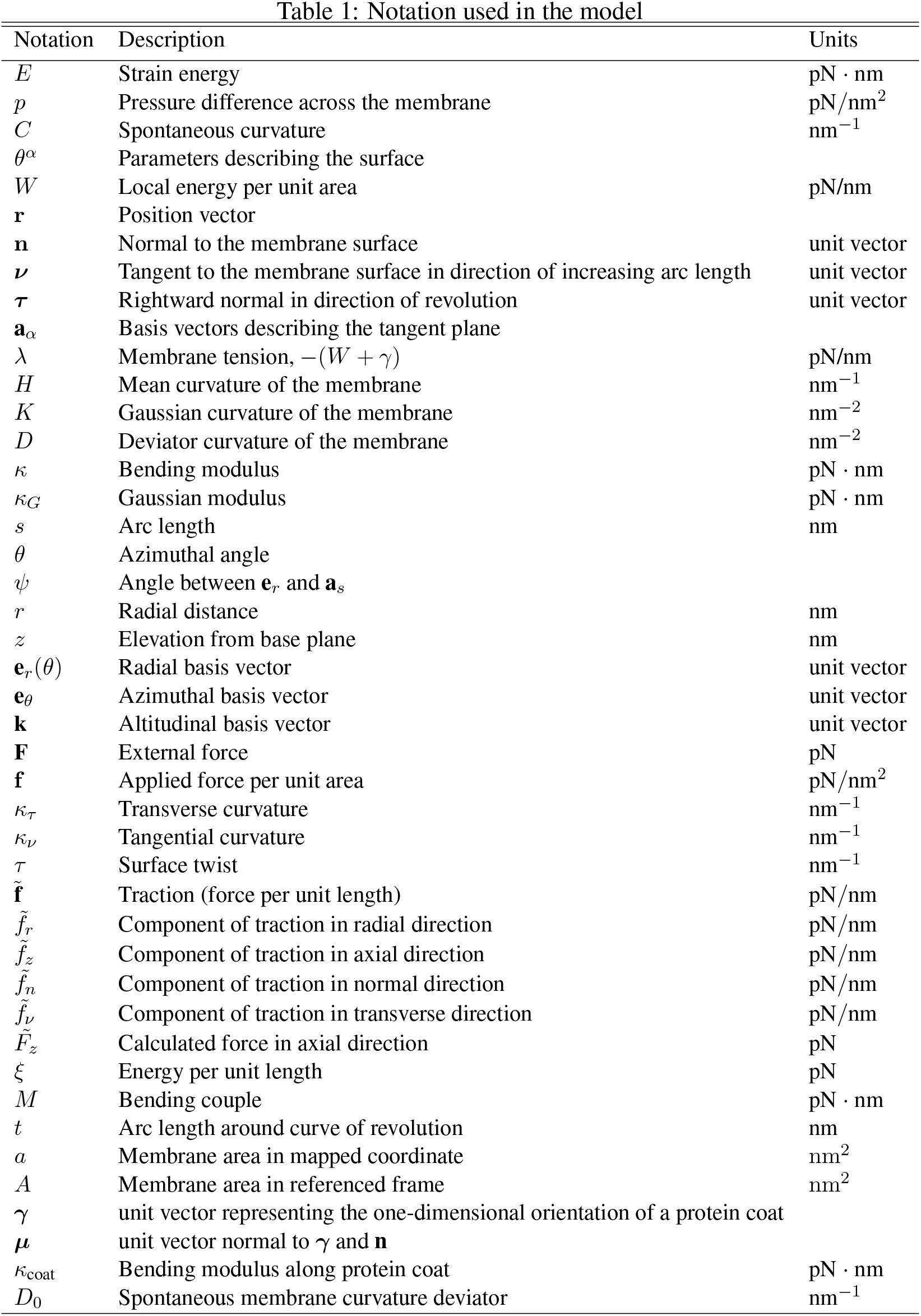
Notation used in the model

| Notation | Description | Units |
| --- | --- | --- |
| $E$ | Strain energy | $\text{pN} \cdot \text{nm}$ |
| $p$ | Pressure difference across the membrane | $\text{pN}/\text{nm}^2$ |
| $C$ | Spontaneous curvature | $\text{nm}^{-1}$ |
| $\theta^\alpha$ | Parameters describing the surface | |
| $W$ | Local energy per unit area | $\text{pN}/\text{nm}$ |
| $\mathbf{r}$ | Position vector | |
| $\mathbf{n}$ | Normal to the membrane surface | unit vector |
| $\boldsymbol{\nu}$ | Tangent to the membrane surface in direction of increasing arc length | unit vector |
| $\boldsymbol{\tau}$ | Rightward normal in direction of revolution | unit vector |
| $\mathbf{a}_\alpha$ | Basis vectors describing the tangent plane | |
| $\lambda$ | Membrane tension, $-(W + \gamma)$ | $\text{pN}/\text{nm}$ |
| $H$ | Mean curvature of the membrane | $\text{nm}^{-1}$ |
| $K$ | Gaussian curvature of the membrane | $\text{nm}^{-2}$ |
| $D$ | Deviator curvature of the membrane | $\text{nm}^{-2}$ |
| $\kappa$ | Bending modulus | $\text{pN} \cdot \text{nm}$ |
| $\kappa_G$ | Gaussian modulus | $\text{pN} \cdot \text{nm}$ |
| $s$ | Arc length | $\text{nm}$ |
| $\theta$ | Azimuthal angle | |
| $\psi$ | Angle between $\mathbf{e}_r$ and $\mathbf{a}_s$ | |
| $r$ | Radial distance | $\text{nm}$ |
| $z$ | Elevation from base plane | $\text{nm}$ |
| $\mathbf{e}_r(\theta)$ | Radial basis vector | unit vector |
| $\mathbf{e}_\theta$ | Azimuthal basis vector | unit vector |
| $\mathbf{k}$ | Altitudinal basis vector | unit vector |
| $\mathbf{F}$ | External force | $\text{pN}$ |
| $\mathbf{f}$ | Applied force per unit area | $\text{pN}/\text{nm}^2$ |
| $\kappa_\tau$ | Transverse curvature | $\text{nm}^{-1}$ |
| $\kappa_\nu$ | Tangential curvature | $\text{nm}^{-1}$ |
| $\tau$ | Surface twist | $\text{nm}^{-1}$ |
| $\tilde{\mathbf{f}}$ | Traction (force per unit length) | $\text{pN}/\text{nm}$ |
| $\tilde{f}_r$ | Component of traction in radial direction | $\text{pN}/\text{nm}$ |
| $\tilde{f}_z$ | Component of traction in axial direction | $\text{pN}/\text{nm}$ |
| $\tilde{f}_n$ | Component of traction in normal direction | $\text{pN}/\text{nm}$ |
| $\tilde{f}_\nu$ | Component of traction in transverse direction | $\text{pN}/\text{nm}$ |
| $\tilde{F}_z$ | Calculated force in axial direction | $\text{pN}$ |
| $\xi$ | Energy per unit length | $\text{pN}$ |
| $M$ | Bending couple | $\text{pN} \cdot \text{nm}$ |
| $t$ | Arc length around curve of revolution | $\text{nm}$ |
| $a$ | Membrane area in mapped coordinate | $\text{nm}^2$ |
| $A$ | Membrane area in referenced frame | $\text{nm}^2$ |
| $\boldsymbol{\gamma}$ | unit vector representing the one-dimensional orientation of a protein coat | |
| $\boldsymbol{\mu}$ | unit vector normal to $\boldsymbol{\gamma}$ and $\mathbf{n}$ | |
| $\kappa_{\text{coat}}$ | Bending modulus along protein coat | $\text{pN} \cdot \text{nm}$ |
| $D_0$ | Spontaneous membrane curvature deviator | $\text{nm}^{-1}$ |

While Eqs. 1 & 3 are general expressions that are independent of coordinates, for illustrative purposes we will restrict further analysis to rotationally symmetric membrane deformations for ease of analysis (Fig. 2). Using principles of force balance one can derive the “shape” equation and the tangential balance equation for the Helfrich energy (see Supplement for detailed derivations). The traction, which is the force per unit length, across any boundary of constant *z* is given by

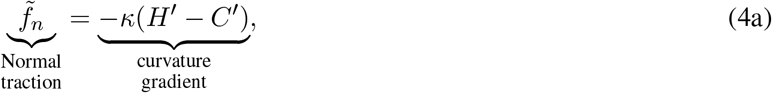

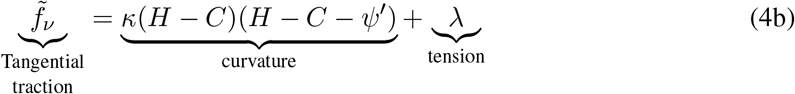

where *ψ* is the angle the membrane makes with the horizontal (see Fig. 2), λ is the local membrane tension, and ()’ denotes a derivative with respect to arc-length s, e.g. *H*’ = d*H*/d*s*.

From the above equations, we see that the normal traction, 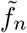, captures the effect of curvature gradients while the tangential traction, 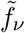, captures the effect of local membrane tension and curvature. A complete derivation of the stress balance and the governing equations of motion is presented in the Supplement. Additional derivations of traction including spatially heterogenous bending and Gaussian moduli, asymptotic approximations for small radius as well as anisotropic spontaneous curvature are presented in the Supplement.

### Interpretation of traction

Traction, which has the units of force per unit length, was initially introduced by physicists as a result of Noether’s theorem (Capovilla and Guven, 2002a; Guven, 2004; Capovilla and Guven, 2004). This theorem states that for any elastic surface that is in equilibrium, there exists a unique traction distribution such that its divergence is conserved (Guven, 2004). Mechanically, the traction distribution gives us information about the response of the membrane to externally applied loading, including forces acting on the membrane or protein-mediated bending. Numerous studies have derived these equations mathematically and sought to explain them in a biophysical context. Capovilla and Guven (Capovilla and Guven, 2002b,a, 2004) invoked the action-reaction law – if one were to cut the membrane along any curve, 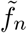 and 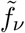 are the forces per unit length of the curve in the normal and tangential directions respectively that the membrane on one side of the cut exerts on the other. Further-more, the expressions for tractions (Eq. 4) reduce to their corresponding fluid analogues for negligible membrane rigidity and pressure difference. Thus, we can interpret the normal and tangential tractions as follows – the tangential traction distribution tracks the gradient in ‘effective’ surface tension (discussed below) while the normal traction distribution contains information regarding a force balance performed normal to the membrane at every point. Further physical interpretations of these quantities can be obtained based on the particular biological phenomena, as illustrated below by examining two fundamental membrane deformations – tubes and buds.

### Axial force and effective line tension

We obtain the formulae for traction in the axial and radial directions obtained by projecting the normal and tangential tractions onto these axes (Eqs. S28) (full derivation in Supplement). We can then calculate the magnitude of an applied axial force on the membrane by integrating the axial component of the traction (Eq. S28b) along the circumference of the bounding curve *∂ω*, yielding

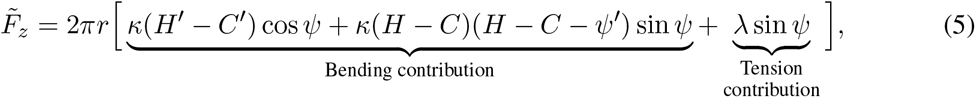

where 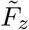 is the axial force generated in response to an external load.

An energy per unit length, *ξ*, associated with deformations in the radial direction can be found by integrating the radial traction along the curve *∂ω* (Fig. 2), as

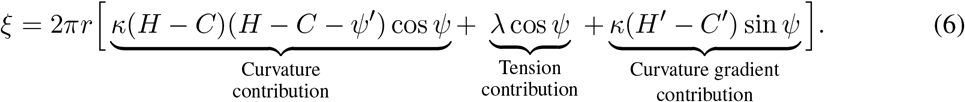

*ξ* can be interpreted as an “effective” line tension (Seifert, 1997). While line tension denotes the force acting at the boundary of two interfaces – e.g. inward force for a liquid droplet on a hydrophobic substrate and an outward force on a hydrophilic substrate (Buehrle et al., 2002; Liu et al., 2006), the “effective” line tension predicts a general resistive force acting at every point opposing any change in the membrane length, regardless of a phase boundary. This ‘force’ is not an actual radial force but represents the change in energy with respect to the characteristic length scale (McDargh et al., 2016); going forward, we refer to it as an energy per unit length.

### Illustrative examples of traction along the membrane

For spherical vesicles, where the mean curvature is constant and in the absence of spontaneous curvature curvature (*C* = 0) and homogeneous composition, the normal traction 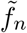 is zero because curvature gradients are zero (Eq. 4a), and the tangential traction, 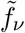, reduces to the membrane tension (λ) (Eq. 4b). This is consistent with previous discussions of membrane tension (Rangamani et al., 2014b). For surfaces with zero mean curvature (minimal surfaces such as catenoids (Powers et al., 2002)) and homogeneous composition, 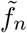 is zero and 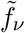 is equal to λ, also consistent with the interpretation of membrane tension for these surfaces (Powers et al., 2002; Chabanon and Rangamani, 2018).

What happens when the mean curvature is not constant or if the membrane is not homogeneous in composition? Given a membrane shape and a constitutive relationship, Eqs. 4a and 4b tell us that we can calculate the local stresses along the membrane. One way of studying shapes is to use images from high resolution microscopy of membrane vesicles of known composition. However, these images can be noisy and obtaining the local curvature and curvature gradients requires fitting the curve with multiple splines or other functions (Lee et al., 2008). Another way to generate membrane shapes is to use simulations. Since our goal is to illustrate the concept of local tractions, we use shapes generated from simulations to elucidate how the normal and tangential tractions are distributed along the membrane. The traction distributions are not the direct output of these simulations; instead they are calculated *a posteriori* using the output shapes from the simulations and the membrane properties, similarly to how one would calculate these distributions from experimentally observed membrane shapes.

### Tether formation due to applied load – revisiting a classical membrane deformation

The formation of membrane tethers in response to a point load is a classic example of force-mediated membrane deformation (Roux et al., 2002; Smith et al., 2004) that has been extensively studied both experimentally (Waugh, 1982; Heinrich et al., 1999) and theoretically (Derényi et al., 2002; Powers et al., 2002; Prévost et al., 2017; Simunovic et al., 2017). This comes as no surprise because a tether is a starting point for understanding membrane deformation in a wide variety of biological contexts including endocytosis, filopodia formation, tubulation in the endoplasmic reticulum, etc. We used this example to validate our method and to identify how normal and tangential tractions contribute to the formation of tethers. We generated a membrane tether by applying a localized force at the pole to mimic a point load, and solved the shape equation for homogeneous bilayers in axisymmetric coordinates (Eq. S17), for a membrane tension of 0.02 pN/nm (simulation details provided in the Supplement).

The normal and tangential traction distributions along the tether are shown in Fig. 3. The absolute value of the normal tractions are highest at the pole as the applied force increases. The membrane curves away from the applied force along the region over which it is applied, and conforms to a stable cylindrical geometry along the rest of the tether and a flat region at the base. The tangential traction has a large positive value along the cylindrical portion of the tether (Fig. 3C) showing that the membrane resists stretching as the tube is pulled out. The tether cap has a negative tangential traction because of the membrane tension heterogeneity (Eq. S10) induced by the application of the load. The corresponding radial and axial traction components (Eqs. S28a, S28b) plotted along the equilibrium shapes are shown in Fig. S1.

**Figure 3:**
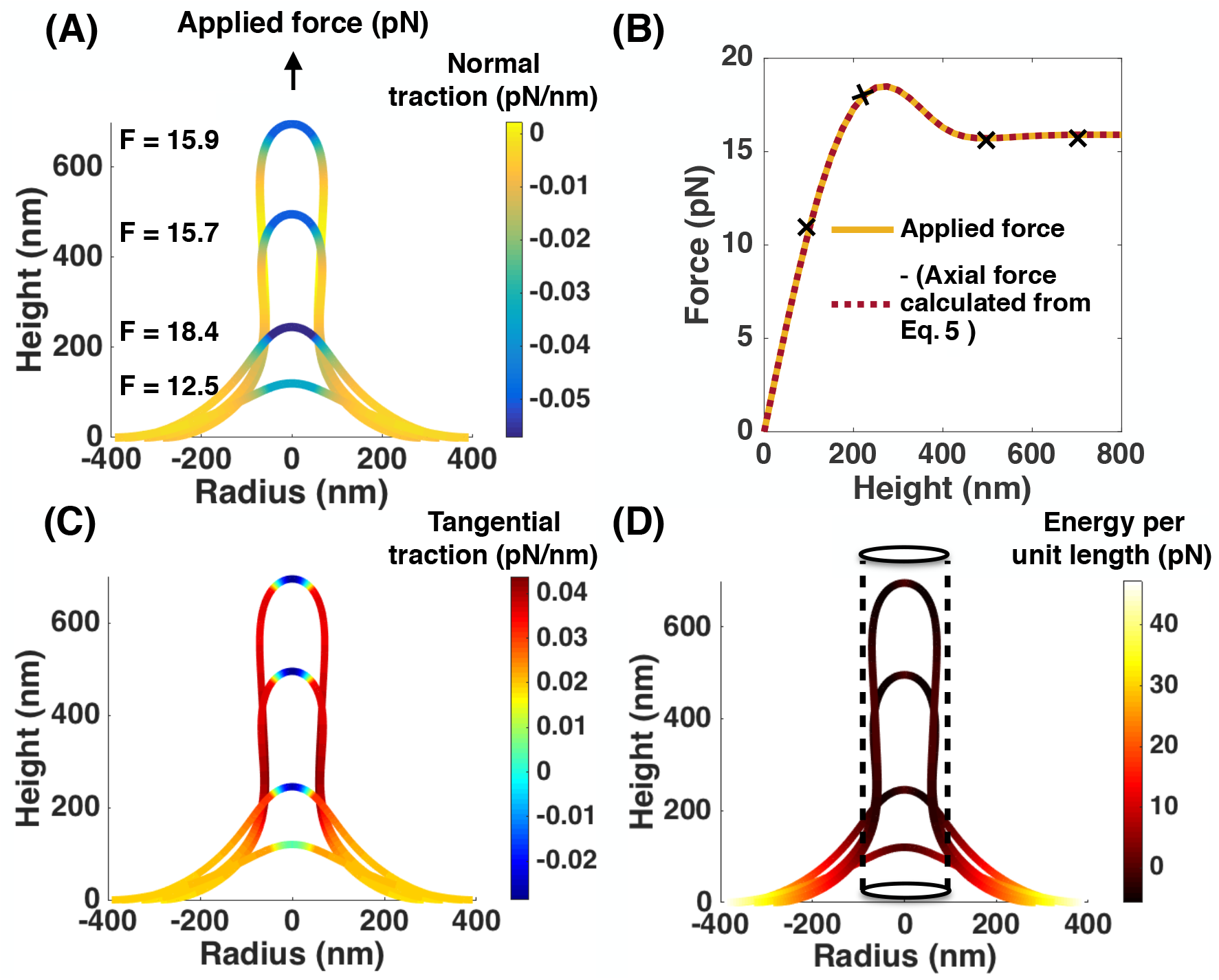
Analysis of normal and tangential traction for membrane tethers. (A) Normal traction distribution along four membrane tether shapes obtained by applying a point load of the specified magnitude at the pole, λ_0_ = 0.02 pN/nm, *κ* = 320 pN · nm. (B) Magnitude of axial force as a function of tether height, showing an exact match between the force (Eq. 5) calculated from the traction distribution and obtained directly from the simulation. (C) Tangential traction distribution along the membrane shapes for conditions shown in (A). (D) Energy per unit length calculated using Eq. 6 along the four membrane shapes shown in (A). The dashed lines outline the equilibrium geometry for a membrane cylinder, 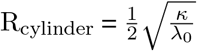.

As expected, the negative of the axial force (Eq. 5), evaluated at the base of the geometry, exactly matches the force-extension relationship for tether formation obtained directly from the simulation (see Fig. 3B), showing that the local stresses along a membrane shape can help us evaluate the applied forces. We also considered the role of a large turgor pressure that opposes the membrane invagination, mimicking the situation in yeast endocytosis. (Basu et al., 2014; Aghamohammadzadeh and Ayscough, 2009; Dmitrieff and Nédélec, 2015). Transmembrane pressure results in an additional term in the axial traction (see Eq. S29). As seen in Figs. S2 and S3, an excellent match between the applied load and the calculated force from the traction distribution is obtained for simulations with pressure by modifying our expression for force. We further verified that our results are independent of the system constraints (i.e. conserved arc length or surface area), confirming that changes in membrane area does not change the validity of our approach (Fig. S7).

What information do the tangential tractions contain? The tangential tractions play an important role in squeezing the membrane neck and holding the cylindrical configuration during membrane elongation (see Fig. S1). Consequently in Fig. 3D, the point of zero ‘effective’ line tension corresponds to the the dotted cylinder, which has a radius of 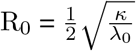 (Derényi et al., 2002). This equilibrium cylinder has no curvature gradient, leading to zero ‘effective’ line tension. The calculated values of energy per unit length inside the cylinder are negative while those outside are positive, suggesting that the “effective” line tension indicates the extent of deviation from the idealized cylindrical geometry. A negative energy per unit length here refers to the fact that there exists a negative radial force at that point (McDargh et al., 2016). Additionally, the value of *ξ* at the neck is ∼ 3pN, providing an estimate of the effective line tension required to form a neck in tethers.

### Traction along tubes is highly dependent on mechanisms of membrane deformation and on resistive force

Do all membrane tubes have the same traction distribution? In order to answer this question, we compared membrane shapes that look superficially similar and calculated the traction profiles along them (Fig. 4). We show that different tubes can have very different tractions depending on the mechanism of membrane deformation and the resisitive forces that are acting on them. We begin by comparing electron micrographs of yeast endocytic invaginations in mutant cells lacking the BAR-domain proteins Bzz1, Rvs167 and wild-type cells (Kishimoto et al., 2011) (Figs. 4A and 4E respectively). Because force from actin assembly is the primary driver of membrane deformation in this process (Kukulski et al., 2012), we assume that the deformation in the mutant cell is a result of having only an applied force at the tip of the invagination (Fig. 4B). In the wild-type, we assume that the BAR domain proteins induce an anisotropic spontaneous curvature locally (e.g. tabulation) (Frost et al., 2009) (Fig. 4F, see Fig. S10 and Supplement for implementation and traction calculation). These assumptions between the mutant and wildtype cells are simplifications, but serve to illustrate the differences in traction distribution. In particular, the tangential traction in the wild-type case (Fig. 4H) is nearly zero near the tip of the bud and highest near the base, in stark contrast to the mutant, which lacks additional curvature generation and therefore is high all along the tube (Fig. 4D). These results suggest that the BAR domain proteins can act as a barrier to the stresses induced by the axial force, which is consistent with recent experimental evidence that points to a potential scission mechanism (Simunovic et al., 2017). Indeed, a negative normal traction at the tube base in Fig. 4G demonstrates a tendency for the neck to shrink in size.

**Figure 4:**
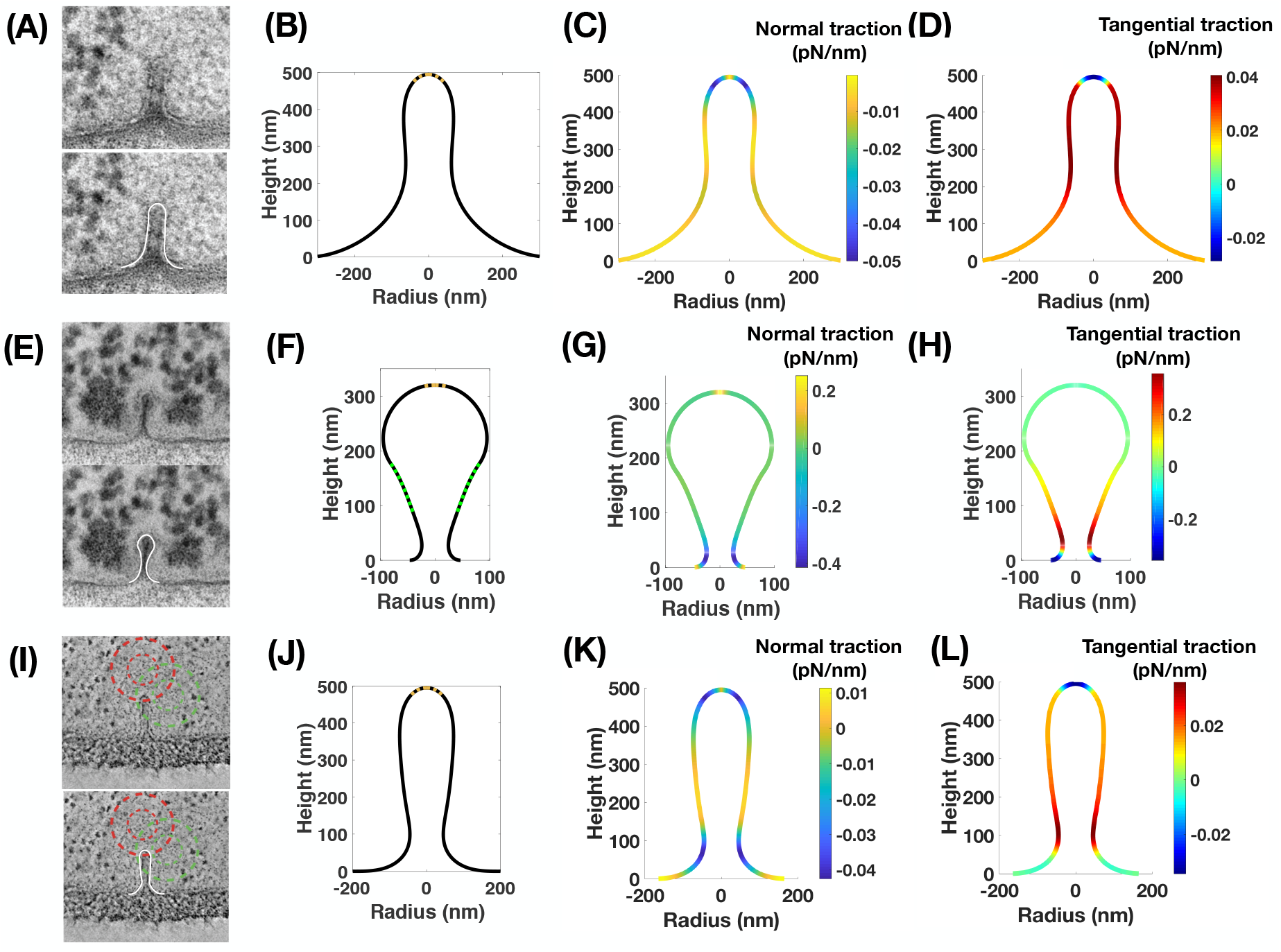
Comparison of normal and tangential tractions between multiple mechanisms of membrane tether formation. (A) EM image of an endocytic PM invagination in a bzz1Δrvs167Δ yeast cell (Kishimoto et al., 2011). *Top inset* – Original EM image, *Bottom inset* – EM image with traced membrane shape (white). (B) Simulation membrane shape obtained by application of a point force (brown), λ_0_ = 0.02 pN/nm, *k* = 320 pN η nm. (C) Normal traction distribution along the membrane shape in (B). (D) Tangential traction distribution along the membrane shape in (B). (E) EM image of an endocytic PM invagination in a wild type (WT) yeast cell (Kishimoto et al., 2011). *Top inset* – Original EM image, *Bottom inset* – EM image with traced membrane shape (white). (F) Simulation membrane shape obtained by application of an anisotropic spontaneous curvature (green) along the tubular section of a membrane tether, λ_0_ = 0.02 pN/nm, *κ* = 320pN η nm, *C* = –0.01nm^−1^, D = 0.01 nm^−1^. (G) Normal traction distribution along the membrane shape in (F). (H) Tangential traction distribtuion along the membrane shape in (F). (I) ET (electron tomography) image of an endocytic invagination in budding yeast (Kukulski et al., 2012). *Top inset* – Original EM image, *Bottom inset* – EM image with traced membrane shape (white). (J) Simulation membrane shape obtained by application of a point force (brown) against an equivalent pressure to the membrane tension in (B), λ_0_= 0pN/nm, *κ* = 320pN *κ* nm, *p* = 0.3kPa. (K) Normal traction distribution along the membrane shape in (J). (L) Tangential traction distribution along the membrane shape in (J).

The previous simulations were conducted using a membrane tension that is applicable to mammalian cells (Sens and Plastino, 2015). However, turgor pressure is thought to be the primary opposing force in yeast endocytosis (Aghamohammadzadeh and Ayscough, 2009). To investigate the role of turgor pressure, we performed a simulation in which the value of the turgor pressure was set such that the radius of the resulting tube (Fig. 4J) would match that of the tube generated using membrane tension (Fig. 4B). The normal traction distribution in this case (Fig. 4K) is strikingly different; there is a large negative normal traction at the base of the tube indicating that turgor pressure acts to induce the formation of a neck. The tangential traction (Fig. 4L) is no longer uniform on the tube and is again greatest just above the narrowing of the tube at the base. Though these simulations are only meant to capture the approximate shapes of the membrane in these different cases and not necessarily match the length scales or parameter values with respect to the biological situations, they serve to illustrate the point that the quantitative differences in the deformations against membrane tension and turgor pressure can be realized by the calculating the local tractions along the membrane shape.

### Formation of buds due to spontaneous curvature is characterized by emergent line tension

Phase separation and lipid domains are classical mechanisms of bud formation and vesiculation (Richmond et al., 2011). Previously, we and others have shown that protein-induced heterogeneity on the membrane can be modeled using a spontaneous curvature field (Steigmann, 1999; Agrawal and Steigmann, 2009b; Rangamani et al., 2014b). We used this framework to investigate the nature of membrane tractions generated during budding due to a spontaneous curvature field. We conducted simulations for a constant area of the spontaneous curvature field A = 10, 000 nm^2^ and varied the extent of spontaneous curvature, C, from 0 to 0.032 nm^−1^ (Fig. 5A). We calculated the value of traction for three distinct shapes – a shallow invagination, a U-shaped bud, and a closed Ω- shaped bud (Fig. 5B-D).

**Figure 5:**
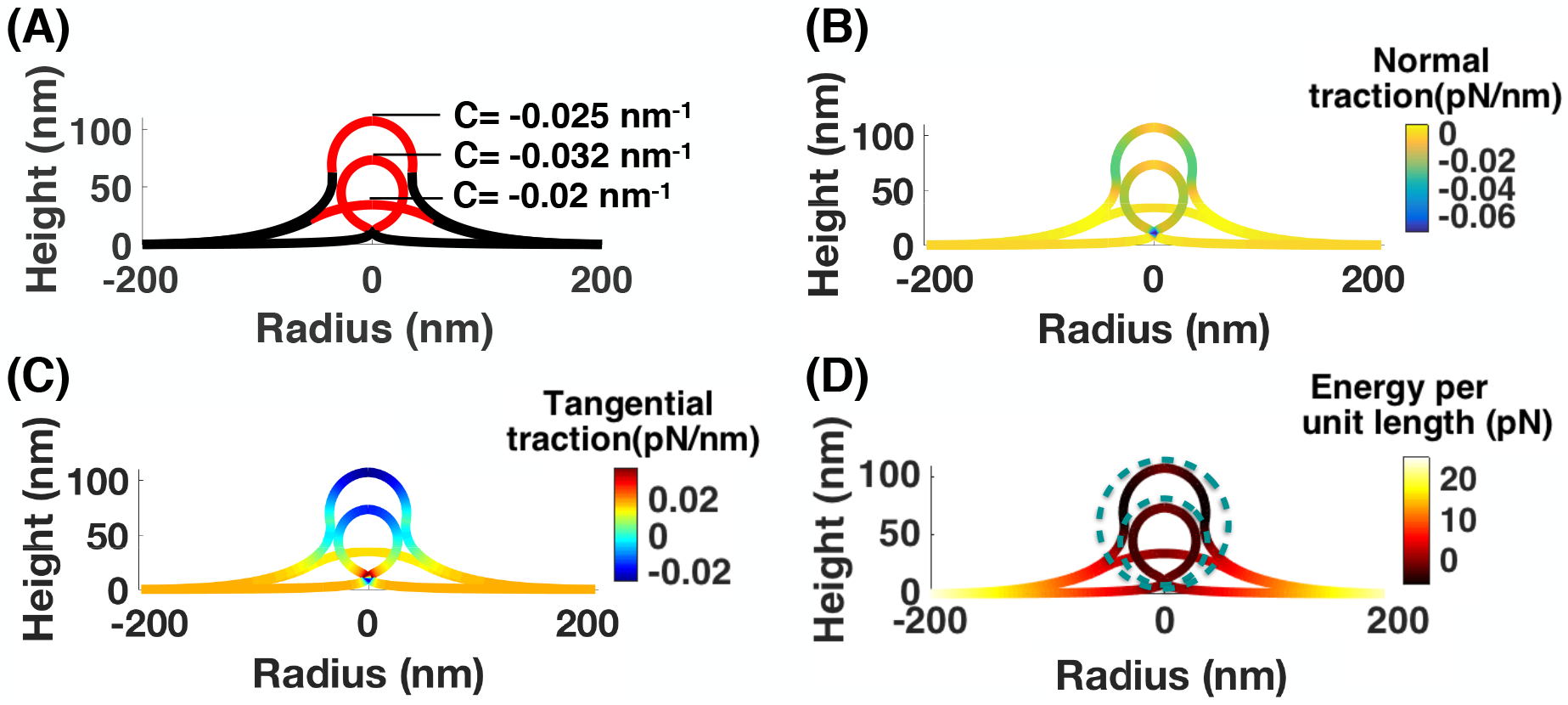
Analysis of budding due to protein-induced spontaneous curvature and calculation of line tension. Simulations were conducted with (A = 10,053 nm^2^) spontaneous curvature at the center of an initially flat patch increasing from *C* = 0 to *C* = 0.032 nm^−1^, λ_0_ = 0.02pN/nm, *κ* = 320 pN η nm, *p* = 0pN/nm^2^ (Hassinger et al., 2017). (A) Membrane shapes for three different spontaneous curvature distributions with the value of C indicated in the red region and zero in the black region. (B) Normal traction along the membrane for the shapes shown in (A). (C) Tangential traction distribution along the shapes shown in (A); (D) Energy per unit length distribution for the three different shapes. The dashed line circles outline spheres with mean curvatures *H* = 0.032 nm^−1^ (smaller circle) and *H* = 0.025 nm^−1^ (larger circle).

The normal traction is negative along the applied spontaneous curvature field indicating a sharper change in mean curvature compared to the applied spontaneous curvature (*H*’ > *C*’ in Eq. 4a). At the neck, where 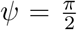, normal traction is maximum and acts purely inward, representing the tendency of the membrane to form small necks. The tangential traction shows a change in sign from positive to negative as the neck radius becomes smaller. This change in sign highlights the critical role of the gradient in tangential traction in the formation of narrow necks (Hassinger et al., 2017) (Figs. 5B-D). The dashed circles represent the equilibrium spherical vesicles calculated by Helfrich energy minimization 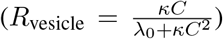 (Hassinger et al., 2017). The positive tangential traction in tent-like small deformations indicates that the membrane resists the bending deformation; however, in the U-shaped and closed buds, the negative tangential traction along the cap acts to pull the membrane inward and favors the adoption of a highly curved shape. The radial and axial tractions distribution along all three shapes are shown in Fig. S4 which reveals that bud formation by spontaneous curvature is purely driven by radial traction while axial traction is zero everywhere.

Each equilibrium bud divides the membrane into two domains – (*i*) the membrane inside the bud with negative energy per unit length that bends to form a bud and (*ii*) the membrane outside the bud with positive energy per unit length that resists such a deformation. Previously, both modeling and experimental studies have shown that in heterogeneous membranes, line tension can be sufficient for scission of endocytic pits (Liu et al., 2006) or the formation of buds in vesicle experiments (Baumgart et al., 2003, 2005). In the case of an applied spontaneous curvature field, the expression of energy per unit length (Eq. S31) can be interpreted as the actual line tension at the interface of the two phases. Through the process of bud formation, line tension undergoes a sign change from positive (acting outward) to negative (acting inward), effectively transitioning from a tension-dominated regime to a curvature-gradient dominated regime (Fig. 6). This transition from positive to negative line tension with increasing value of spontaneous curvature is also observed in other studies (Dan and Safran, 1998). The value of the energy per unit length at the interface varies between −5 pN to 5 pN, which is of the order of the reported interfacial line tension between coexisting phases in lipid bilayers (Lipowsky, 1992; Liu et al., 2006).

**Figure 6:**
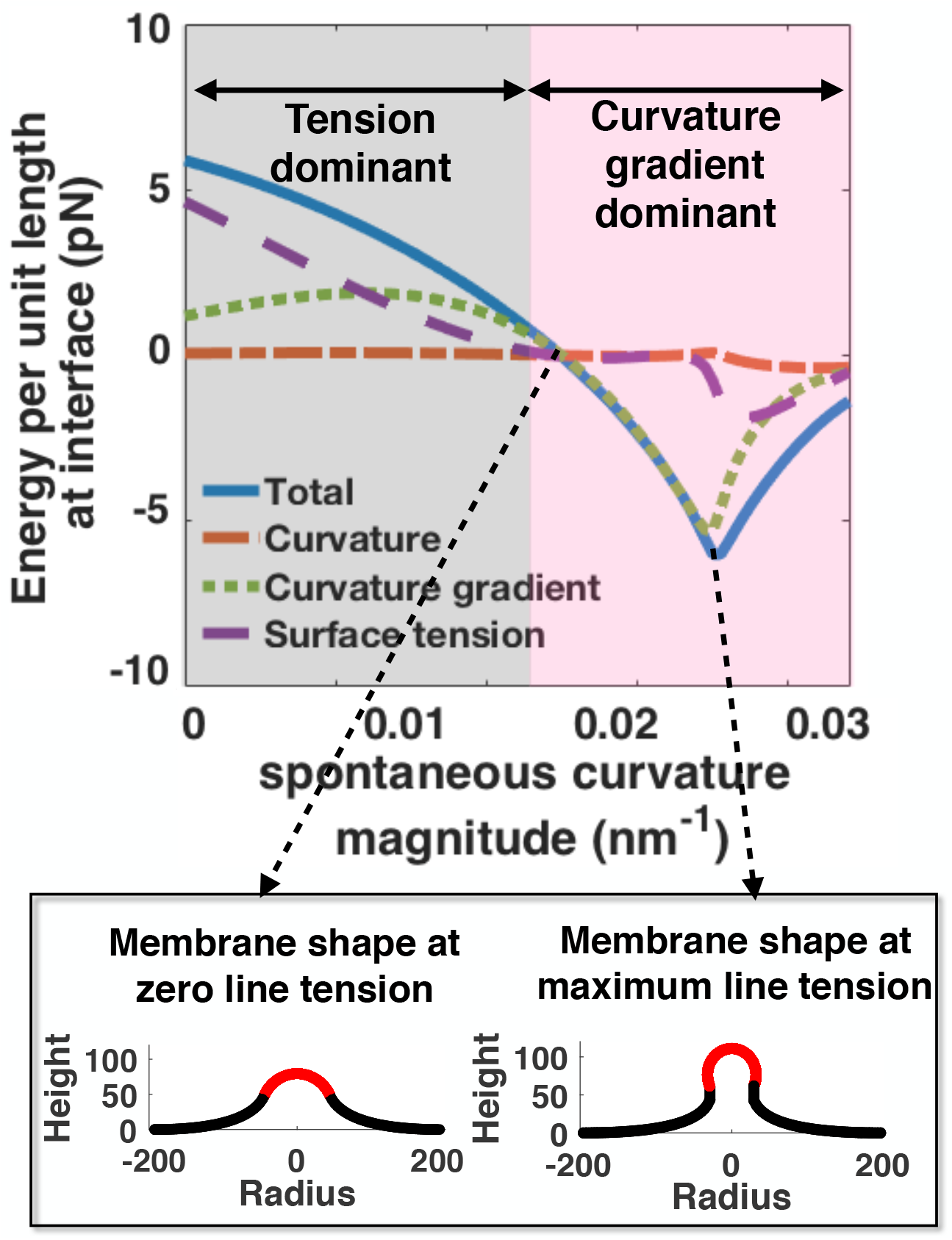
Change in energy per unit length and its components at the interface with increasing spontaneous curvature. Two regimes are observed: a surface tension-dominated regime for small values of spontaneous curvature and a curvature gradient-dominated regime for large vales of spontaneous curvature. The membrane configurations are shown for two spontaneous curvature C = –0.02 nm^−1^, where energy per unit length at interface is zero and *C* = –0.025 nm^−1^, where energy per unit length is maximum. The red domains show the region of spontaneous curvature for the corresponding shapes.

There are two other factors that could affect the traction distribution along the bud – (i) a change in area of the membrane during budding and (ii) spatial heterogeneity in membrane moduli. To explore how the change of membrane area influences bud formation mediated by protein-induced spontaneous curvature, we conducted a simulation with a fixed available arc-length instead of area (Fig. S6). Similar to the case of a homogenous membrane with fixed area, the energy per unit length at the interface changes sign from positive to negative in a range of −5 pN to 5 pN. However, protein segregation on the membrane can lead to heterogeneity in material properties such as bending moduli (Jin et al., 2006). In order to investigate the effect of this spatial heterogeneity in the bending moduli along the membrane surface, we repeated the budding simulation from Fig. 5, assuming that the bending rigidity along the spontaneous curvature field is 7.5 times larger than the bending rigidity of the bare membrane (Fig. S5) (Jin et al., 2006). Because the membrane is stiffer and harder to bend, a wider neck is formed at C = −0.032 nm^−1^ compared to the case of a uniform membrane (Fig. S5A) (Hassinger et al., 2017). This membrane resistance to deformation is observed as a uniform positive normal traction everywhere along the membrane (Fig. S5A). To compare the behavior of the line tension at the edge of the spontaneous curvature field, we ran the budding simulation with the spatially heterogeneous bending moduli up to a larger value of spontaneous curvature (C = −0.035 nm^−1^), in order to have the same range of neck radii as the uniform membrane (Fig. 5E). We can see that the trend of line tension variation versus the spontaneous curvature is almost the same in both cases (Fig. S5E), changing sign from positive to negative followed by a critical point indicating the transition from a U to an Ω-shaped bud. However, the magnitude of line tension is different in the two cases. For small magnitudes of spontaneous curvature (tent shaped buds), the average difference in line tension is ∼ 1 pN. But for large magnitudes of spontaneous curvature (C ≥ −0.0275 nm^−1^, Ω shaped buds), the average line tension for a rigid coat is ∼ 4 pN larger than the line tension in a homogeneous membrane. This larger value of line tension in a heterogeneous membrane has been reported in various experimental measurements (Lipowsky, 1992; Tian et al., 2007), and other theoretical studies (Kuzmin et al., 2005; Semrau and Schmidt, 2009).

### Traction distribution is a signature of distinct budding mechanisms

Conceptually, there are two primary means by which membrane buds can be maintained: an accumulation of protein or lipid-induced spontaneous curvature favoring a spherical geometry, or a constriction force that pinches the membrane into a budded shape. In Fig. 7, we illustrate the traction distribution in these two cases. The upper row represents spontaneous curvature-induced budding, meant to resemble vesicle coat protein (such as the coatomer COPII) mediated budding from the endoplasmic reticulum ((Robinson et al., 2015), Fig. 7A) and the lower row represents budding due to a local constriction force via a contractile ring in budding yeast ((Mozdy et al., 2000), Fig. 7E). Although the two simulated shapes are superficially similar, the traction distributions are quite different. The normal traction distribution for spontaneous curvature budding (Fig. 7C) is similar to the one seen in Fig. 5 where there is a large negative traction at the bud neck, indicating forces acting to minimize the neck radius. Conversely, for the constriction force budding, the normal traction is highly positive at the neck (Fig. 7G), indicating a resistance by the membrane to the applied force. The tangential tractions (Fig. 7D and H) are also quite different. For example, moving from the top to the bottom of the vesicle, the tangential traction in the case of the protein-induced spontaneous curvature budding is initially negative and then positive after the neck (Fig. 7D). However, for the constriction force mediated budding, the tangential traction is positive at first and then negative after the neck (Fig. 7D). This difference in the gradient of tangential traction at the membrane neck serves as a signature for spontaneous curvature mediated vs force mediated bud formation. Thus, the mechanism of curvature generation can be related to the computed traction profile, and some *a priori* knowledge can help uncover these differences (see Figs. 4 and 7).

**Figure 7:**
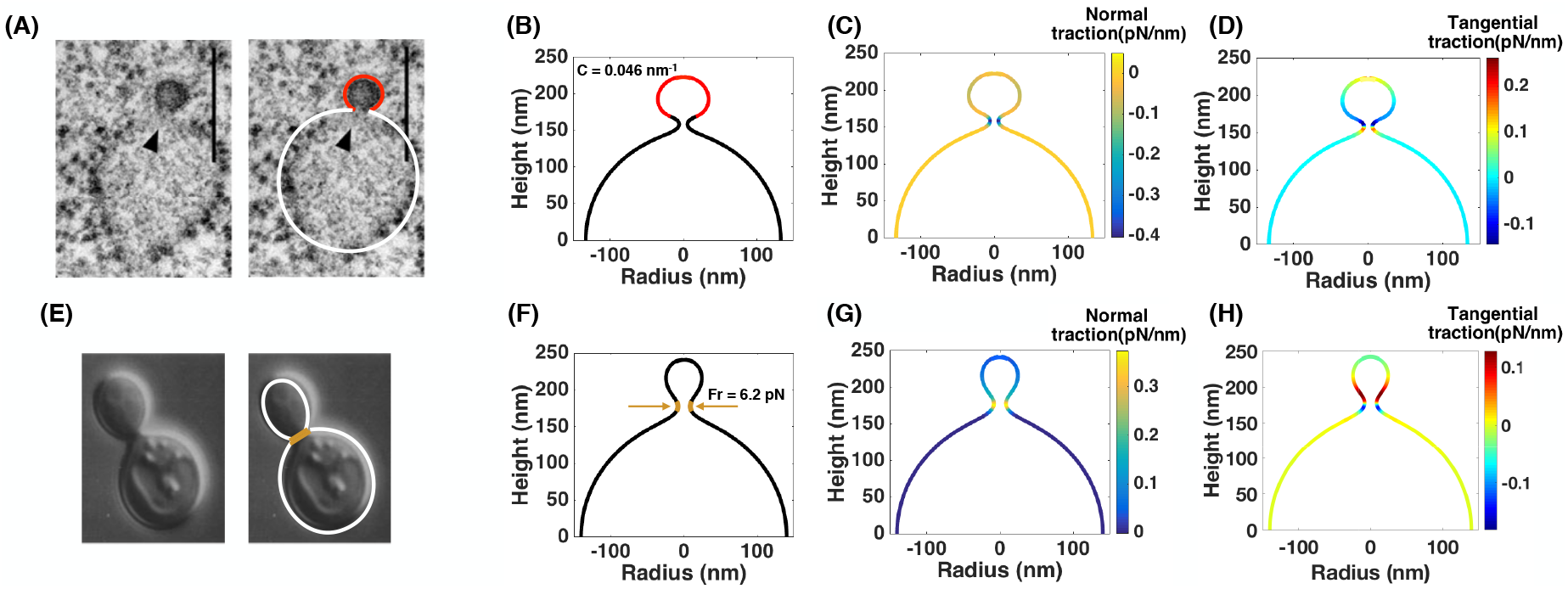
Comparison of normal and tangential tractions between two different mechanisms of membrane budding. (A) EM image of COPII budding from the endoplasmic reticulum (ER) in green algae (Robinson et al., 2015). *Left inset* – Original EM image, *Right inset* – EM image with traced membrane shape. Red - COPII coat, white - bare membrane (B) Simulation of bud formation on a hemispherical cap using a constant spontaneous curvature (C= −0.046 nm^−1^, red) (C) Normal traction distribution along the membrane shape in (B). A large negative normal traction can be seen at the neck of the formed vesicle. (D) Tangential traction distribution along the membrane shape in (B). There is a change in the sign of the tangential traction before and after the bud neck. (E) Brightfield microscopy image of a budding yeast (Mozdy et al., 2000). *Left inset* – Original EM image, *Right inset* – EM image with traced membrane shape. brown - contractile ring at the bud neck. (F) Simulation of bud formation on a hemispherical cap with a constant radial force (F_r_= 6.2pN, yellow) that locally constricts the hemisphere to form a bud. (G) Normal traction distribution along the membrane shape in (F). There is a positive normal traction at the vesicle neck in response to the applied force. (H) Tangential traction distribution along the membrane shape in (F).

Another mechanism of maintaining membrane buds (specific to endocytosis) is through actin- mediated forces where an actin network polymerizes in a ring at the base of the plasma membrane (PM) invagination and is connected to the coat, driving inward movement (Picco et al., 2015; Walani et al., 2015). We have previously considered these cytoskeletal effects in (Hassinger et al., 2017) and show here that the applied forces can be matched to axial forces calculated from traction (Figs. S8, S9) for two orientations of the applied force.

### Sensitivity analysis and sources of errors

In principle, calculating force from shape is at the heart of stress-strain relationships. However, there are some fundamental challenges associated with sources of errors in such a calculation. There are two main sources of errors – error in the measurement of material properties (membrane bending modulus and membrane tension), and error in the measurement of shape. We present some simple analysis of these sources of error in what follows.

### Parametric sensitivity analysis of material properties

Ideally, one would like to define a sensitivity index similar to the parametric sensitivity conducted for systems of chemical reactions, where the sensitivity of a quantity *F_i_* with respect to a parameter *k_j_* is given by 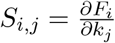 (Varma et al., 2005). However, since we wish to simultaneously explore the effect of both the bending modulus and tension, we use a simple linear calculation of error. Uncertainties in either of these quantities will result in an uncertainty in the traction as well as the calculated axial force and energy per unit length (Eqs. 5 and 6) Here, we assume that the bending modulus and membrane tension can be written as *κ* = *κ*_mean_ ± *κ*_error_ and λ = λ_mean_ ± λ_error_ respectively. Then, by virtue of the relationships in Eqs. 5 and 6, we can estimate the error in the axial force and the energy per unit length as

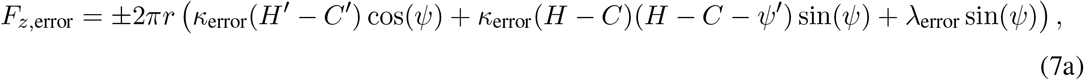

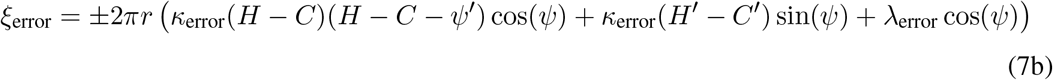

These equations allow us to interrogate how errors in both membrane moduli and membrane tension affect the error in forces. We took our control to be the output of tubulation and budding simulations described in Figs. 3 and 5, respectively. Then, we conducted the same simulations over a range of bending moduli and membrane tensions to reflect a range in error of these two quantities. From these simulations, we (i) calculated the applied force using Eq. 5 for the tube pulling simulations at the peak of the force displacement plot and (ii) the energy per unit length at the phase boundary using Eq. 6 for the budding simulations at the same value of spontaneous curvature. Fig. 8A and 8C show the result of this procedure for a force and energy per unit length respectively that have been normalized to the output from the initial simulations (as indicated by X.) As expected, separately varying either bending modulus or membrane tension is translated into an error in the force and energy per unit length, though the magnitude of the final error does not match that of the input error due to the coupling to shape (Eq. 5 and 6). Next, we investigate the nonlinear effect of varying bending modulus and membrane tension simultaneously on the computed errors. Interestingly, we see that in some cases the error in one parameter is compensated for by the error in the other, as highlighted by the dashed lines which indicate a band of less than 10% total error. This is due to the intrinsic scaling in both tubulation (Derényi et al., 2002; Dmitrieff and Nédélec, 2015) and budding (Hassinger et al., 2017) with respect to bending modulus and membrane tension. Overall, we observe that the final error is not simply a sum of the errors in the two material properties and compensatory behaviours can result (Eq. 7, Fig. 8A, C).

**Figure 8:**
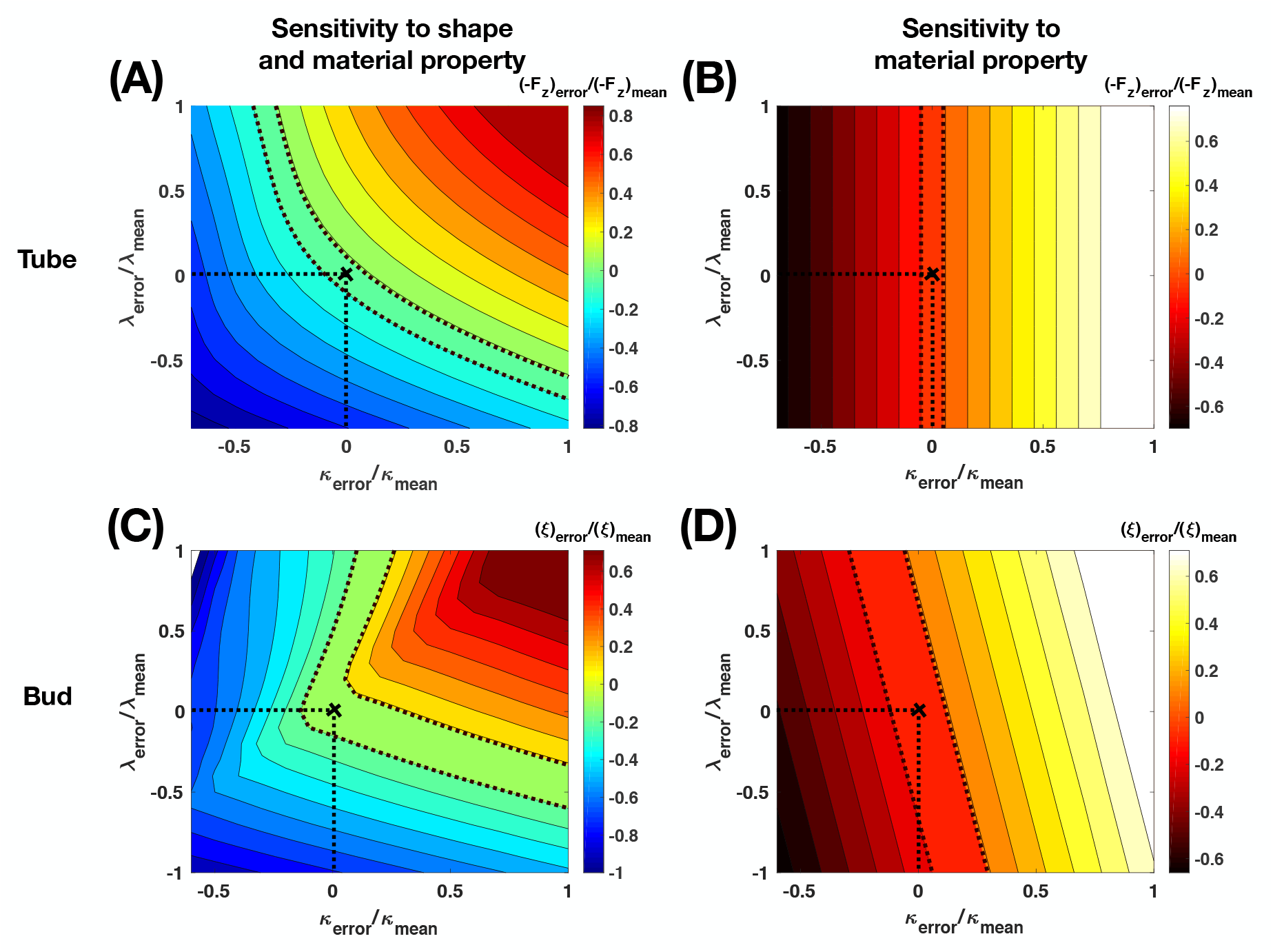
Parametric sensitivity analysis to material properties. Axial force (Eq. 5) and energy per unit length (Eq. 6) were calculated for a variation in the bending rigidity *κ* and membrane tension λ_0_ both in membrane tubes (A-B) and buds (C-D). Dashed lines indicate 10 % error. λ_mean_= 0.02pN/nm, *κ*_mean_ = 320 pN η nm, – (*F_z_*)_mean_ = 18.0167pN (corresponding to a tube of height 300 nm in Fig. 3), *ξ*_mean_ – 6.13pN (corresponding to a spontaneous curvature of 0.0276 nm^−1^ in Fig. 5). The sensitivity analysis was performed in two ways – *(i).* Sensitivity to shape and material property by running multiple simulations corresponding to the different parameter values (A, C) followed by an error calculation with respect to the mean value, *(ii).* Sensitivity to only material property by using a range of parameter values during calculation of axial force (Eq. 5) and energy per unit length (Eq. 6) for a single simulation (mean). (A) Sensitivity to shape and material property in a membrane tube. (B) Sensitivity to only material property in a membrane tube. (C) Sensitivity to shape and material property in a membrane bud. (D) Sensitivity to only material property in a membrane bud.

In the previous calculation, when the membrane modulus and tension were varied, both the characteristic length of the membrane and its shape were affected. We conducted another analysis, where the shape of the membrane was fixed to the control and an error was introduced in the values of bending modulus and membrane tension during the calculation of tractions (Figs. 8B and D). Interestingly, we found that the error in the axial force is independent of the error in membrane tension (Fig. 8B). This is a consequence of calculating the axial force at a point at the base of the deformation where the angle *ψ* is almost zero and so, the tension term contributes less. If one were to instead perform the force balance at a point on the membrane where *ψ* is not zero, the error in the force would again depend on the error in both bending modulus and tension (Fig. S11). This, in principle could be beneficial in the sense that one could minimize the error in determining the axial force by evaluating it at a location where the total error is minimized (e.g. if uncertainty in membrane tension is large, calculate the applied force at the base of the invagination since the calculation is insensitive to error in membrane tension at this location).

In contrast, the phase boundary is located at a particular position on the membrane curve and so must be evaluated at that point. We observe that the dependence of the error in the energy per unit length on bending modulus and membrane tension is no longer non-linear (Fig. 8D) as we fix the shape of the membrane and vary the material properties. Further, we see that the primary dependence of the error in the energy per unit length comes from the error in the bending modulus. Finally, we once again see that the total error is less than the sum of its two contributions due to the coupling to the local membrane shape, as expected from Eq. 7.

### Errors in quantification of shape metrics

One of the largest source of errors in calculating forces arises from imaging modalities for shape itself. Uncertainty in the shape of the membrane will depend on the method used to extract shapes from microscopy images. Additionally, the high curvatures at endocytic sites means that a higher imaging resolution is required. Live-cell light microscopy is limited in resolution (even in superresolution methods (Wäldchen et al., 2015; Sydor et al., 2015)), and traditional electron microscopy following chemical fixation may not fully preserve the shape of the bilayer (Bozzola and Russell, 1999; Stephens and Allan, 2003). To this end, cryo-electron tomography may provide the best preservation, but it suffers from anisotropic resolution as a result of the “missing wedge” effect (Lučić et al., 2013). As a result, error can be introduced into the fundamental position and geometric variables of the constitutive equations associated with the membrane deformation. Errors in the position and shape coordinates, coupled with non-axisymmetric geometries can result in non-linear error propagation in the calculations and their effects are not yet understood.

## Discussion

In this study, we presented a framework for the calculation of axial and radial tractions for non-linear deformations of the membrane in the absence and in the presence of heterogeneities, solely based on the membrane geometry and material properties. From these calculations, we summarize that (a) tether formation requires both axial and radial tractions (Fig. 3) and (b) line tension can be calculated between two phases as an energy per unit length (Fig. 6). Importantly, using different examples of critical membrane shapes that occur in endocytosis and exocytosis, we have demonstrated that the local tractions are directly related to deviations from idealized geometries and can be generated by membrane heterogeneity. Moving forward, this procedure can be useful for the analysis of forces acting on membranes, both in reconstituted systems and in cells.

Using the analysis presented here and having some knowledge of the shape and material properties will allow us to estimate the local stresses acting on the membrane. It is important to note that the tractions calculated here depend on the knowledge of the membrane strain energy and the material properties.

It has been demonstrated that PEGylation of lipids (Lee and Pastor, 2011), amphiphilic block copolymers (Lim et al., 2017), and protein crowding (Snead et al., 2017) can curve and even induce scission of artificial lipid bilayers. Additional energy terms such as adhesion energy, entropic contributions from proteins, protein crowding, tilt, and cytoskeletal interactions will alter the expressions for tractions and introduce more material properties (Rangamani et al., 2014a; Snead et al., 2017; Carlsson, 2018). We also demonstrate that the knowledge of the underlying biophysical mechanism becomes important because the shape of the membrane, particularly in cells, is a many-to-one function (multiple processes can give rise to a similar shape). However, the fundamental principle that shape contains information about the underlying forces will apply regardless of the exact form of the energy used to perform the analysis.

There can be multiple sources of error in the quantification of forces – error in the measurement of material properties, errors in the measurement of the shape itself due to imaging, and finally error in the assumptions about stress-strain relationships themselves. While many of the measurements of material properties are conducted *in vitro*, recently, some studies have begun to measure the *in vivo* structure of lipids and their material properties (Nickels et al., 2017). Interestingly, recent works also suggest that there is no long range propagation of membrane tension in cells, seemingly reducing the uncertainty in calculating tension (Shi et al., 2018). Additionally, efforts will need to focus on the development of image analysis methods to extract the shape of the membrane while reducing noise. There are already quite a few efforts in this direction, although these are focused on tension-based mechanisms in epithelial sheets. Curvature-dependent effects are harder to discern from imaging data (Brodland et al., 2014; Veldhuis et al., 2015). There is also a need for the development of algorithms that do not *a priori* assume symmetry of the shape and can handle irregular geometries. Then, imaging data, which are abundant in the literature (Frost et al., 2009; Dannhauser and Ungewickell, 2012; Snead et al., 2017), can potentially be analyzed and used to populate a database/machine-learning framework. This can then be extended to analyze the shapes of complex structures in cells, which likely include contributions from multiple mechanisms. Finally, an assumption that we have made in this study is to neglect the surrounding fluid flow or inertial dynamics and assume that the membrane is at mechanical equilibrium at all times (Steigmann et al., 2003; Naghdi, 1973; Deserno, 2015). This assumption is commonly used in the modeling of membrane curvature to keep the mathematics tractable (Steigmann, 1999; Deserno, 2015). While the Helfrich model has been used by us and others with great success, the role of these dynamics of deformations, thermal fluctuations (Monzel and Sengupta, 2016; Steinkühler et al., 2018), and multiscale models will be needed to truly appreciate different spatial and temporal scales of forces. As a small step in this direction, we have implemented a modified form of the Helfrich energy including deviatoric effects to consider the anisotropic nature of spontaneous curvature (Fig. S10). While our current focus has been on explaining the mathematical and physical basis of local tractions and how these tractions can be used to understand important experimental systems and biological processes, to close the gap between modeling and experiments, future efforts will need to focus on relaxing the assumption of rotational symmetry and the ability to estimate local tractions in experimentally observed membrane shapes.

**Table 2:**
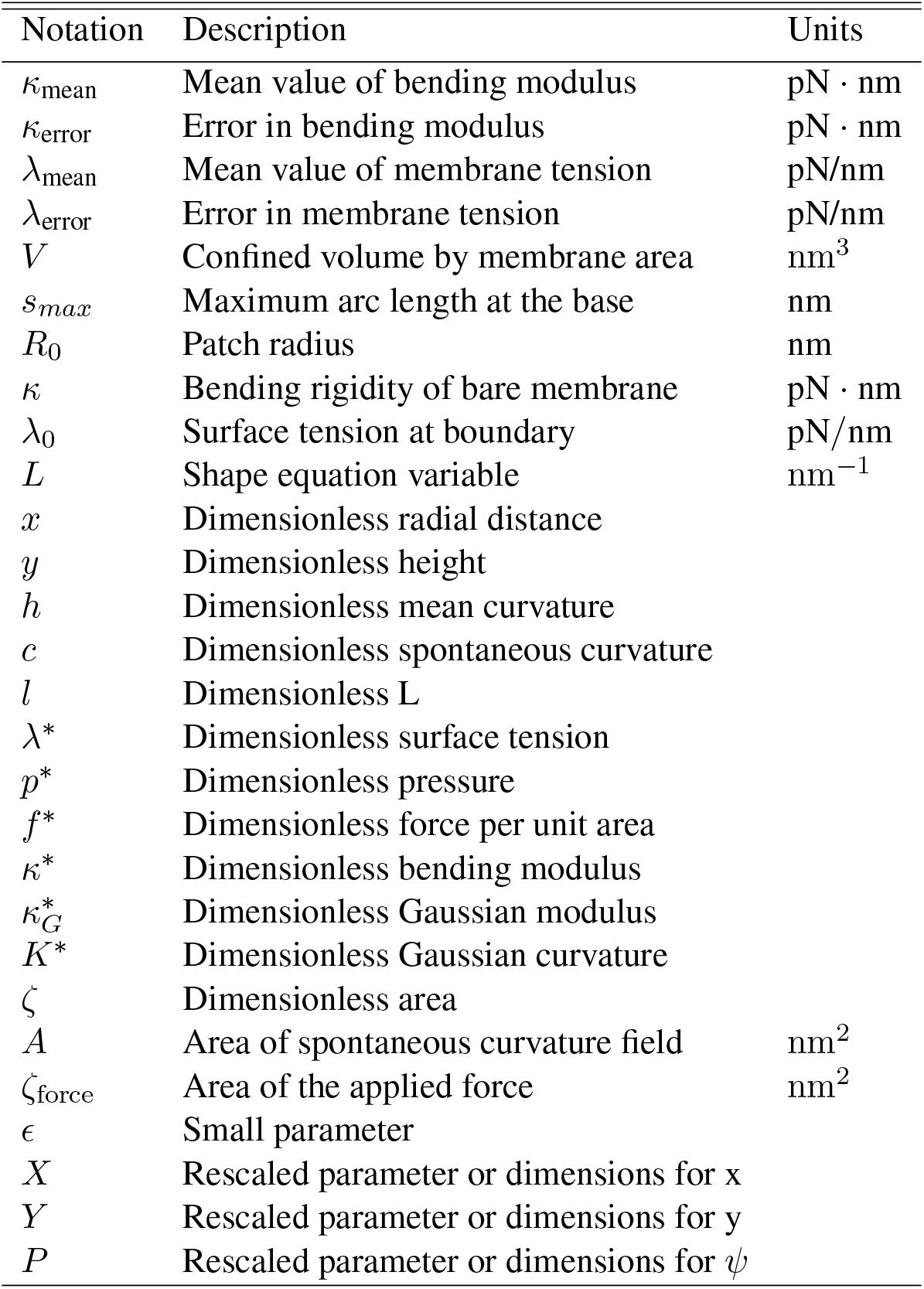
Notation used in the model

## Acknowledgments

This work was supported by ARO W911NF-16-1-0411, AFOSR FA9550-15- 1-0124, and NSF PHY-1505017 grants to P.R. J.C.S. was supported by NIH R01GM112065. R.V. was supported by the UCSD Frontiers of Innovation Scholars Program (FISP) G3020. H.A. was supported by a fellowship from the Virtual Molecular Cell Consortium (VMCC), a program between UCSD and the Scripps Research Institute. J.E.H. was supported by the Department of Defense (DoD) through the National Defense Science & Engineering Graduate Fellowship (NDSEG) Program. The authors would also like to thank Prof. George Oster and Prof. David Steigmann for initial discussions and Drs. Morgan Chabanon and Matthew S. Akamatsu for their suggestions on improving the manuscript.

## 1 Table of notation

## 2 Model development

### 2.1 Assumptions

- Membrane curvature generated due to forces or protein-induced spontaneous curvature is much larger than the thickness of the bilayer. Based on this assumption, we model the lipid bilayer as a thin elastic shell with a bending energy given by the Helfrich-Canham energy, which is valid for radii of curvatures much larger than the thickness of the bilayer [1, 2].
- We neglect the surrounding fluid flow or inertial dynamics and assume that the membrane is in mechanical equilibrium at all times [3]. This assumption is commonly used in the modeling of membrane curvature to keep the mathematics tractable [4].
- The membrane is incompressible because the energetic cost of stretching the membrane is high [5]. This constraint is implemented using a Lagrange multiplier [6, 7].
- Finally, for simplicity in the numerical simulations, we assume that the membrane in the region of interest is rotationally symmetric (Fig. 2).

### 2.2 Equations of motion

At equilibrium, the integration of local energy density (W) over the total membrane surface area ω gives the strain energy of the system written as [8–10]

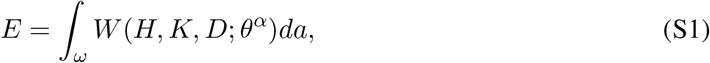

where E is total strain energy, H is the mean curvature of the surface, K is the Gaussian curvature, D is the curvature deviator, and *θ^α^*{*α* = 1,2} denotes the surface coordinate.

To impose the area incompresibility condition, we can rewrite the energy equation Eq. S1 using a Lagrange multiplier

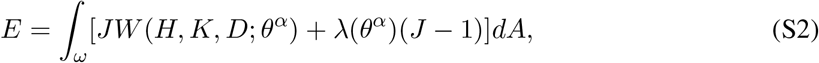

where λ is a Lagrange multiplier and

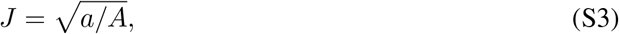

is the local areal stretch due to mapping from a reference frame to the actual surface.

Minimization of the energy Eq. S2 by usage of the variational approach gives the governing shape equation and the incompressibility condition in a heterogeneous membrane

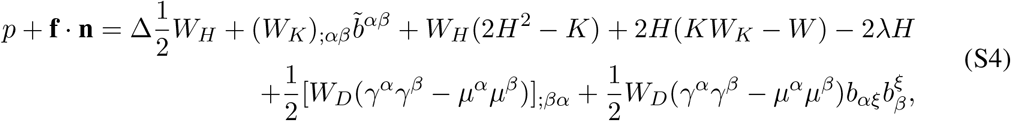

and

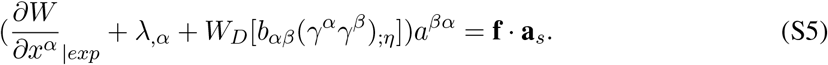

where Δ(·) is the surface Laplacian, p is the pressure difference across the membrane, **f** is any externally applied force per unit area on the membrane, **n** is the normal vector to the surface, **a**_*s*_ is the tangent vector on surface, *a*^*αβ*^ is the dual metric, *b*_*αβ*_ are the coefficients of the second fundamental form, 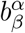 are the mixed components of the curvature, 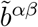 is the co-factor of the curvature tensor, and ()_|*exp*_ denotes the explicit derivative with respect to coordinate *θ^α^*. Also, *γ^α^* and *μ^α^* are the projections of γ and *μ* along the tangential vectors with

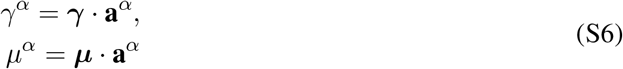

where γ is a unit vector representing the orientation of a protein coat tangential to the one-dimensional curve on the surface, and *μ* is a unit vector defined by

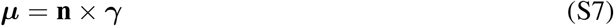

In what follows, we explore different commonly used forms of energy as follows

i. Helfrich energy for isotropic spontaneous curvature
ii. Helfrich energy for anisotropic spontaneous curvature

#### 2.2.1 Helfrich energy for isotropic spontaneous curvature

For a lipid bilayer membrane, we use a modified version of the Helfrich energy to account for the spatial variation of spontaneous curvature [7, 11, 12],

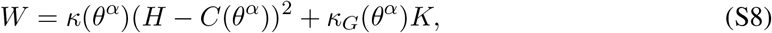

C is the spontaneous curvature, and *κ* and *κ_G_* are bending and Gaussian modulii respectively, which in general case of heterogeneous membrane can vary along thesurface coordinate. It should be mentioned that Eq. S8 is different from the standard Helfrich energy by a factor of 2. We take this net effect into consideration by choosing the value of the bending modulus to be twice that of the standard value of bending modulus typically used for lipid bilayers [1].

Substituting the Helfrich energy function Eq. S8, in Eqs. S4 and S5

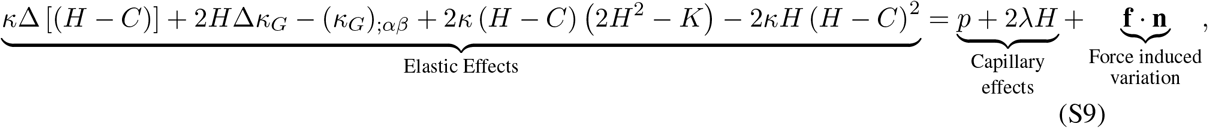

and

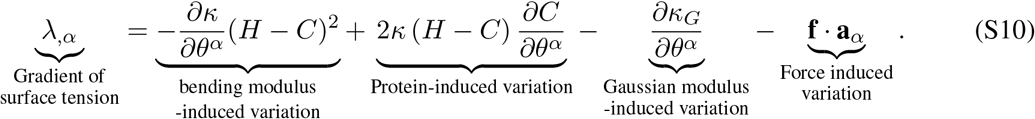

#### 2.2.2 Helfrich energy for an anisotropic curvature

In order to represent anisotropic curvature generated due to membrane-proteins interactions, we consider the local energy density function as [9, 10]

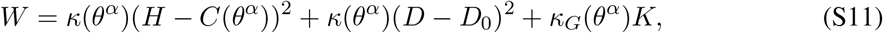

where *D*_0_ is spontaneous membrane curvature deviator. Substituting this form of energy density Eq. S11 in Eqs. S4 and S5 give

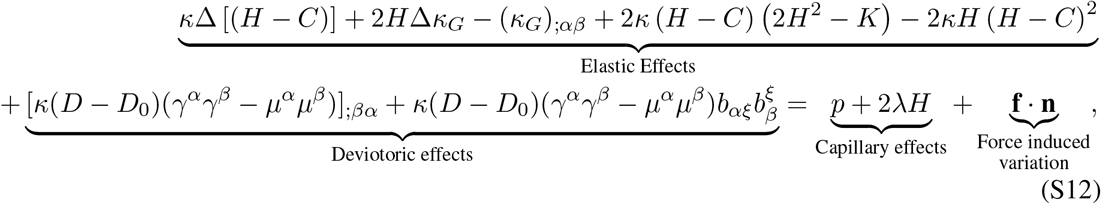

and

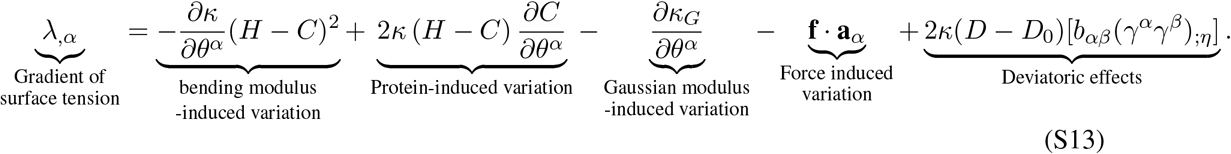

### 2.3 Axisymmetric coordinates

#### 2.3.1 Equations of motion for isotropic curvature

We parameterize a surface of revolution (Fig. 1B) by

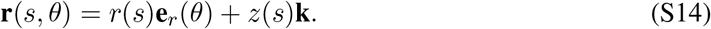

We define *ψ* as the angle made by the tangent with respect to the horizontal. This gives *r*’(*s*) = cos(*ψ*), *z*’(*s*) = sin(*ψ*), which satisfies the identity (*r*’)^2^ + (*z*’)^2^ = 1. Using this, we define the normal to the surface as **n** = –sin *ψ***e**_*r*_ (*θ*) + cos *ψ***k**, the tangent to the surface in the direction of increasing arc length as **ν** = cos *ψ***e**_*r*_(*θ*) + sin*ψ***k**, and unit vector ***τ*** = **e**_*θ*_ tangent to the boundary *∂ω* in the direction of the surface of revolution (see Fig. 1B).

This parameterization yields the following expressions for tangential (*κ_ν_*) and transverse (*κ_τ_*) curvatures, and twist (*τ*):

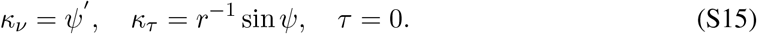

The mean curvature (*H*) and Gaussian curvature (*K*) are obtained by summation and multiplication of the tangential and transverse curvatures

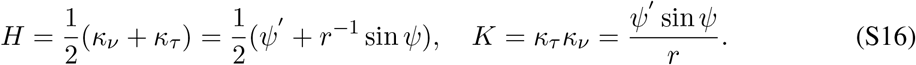

Defining 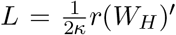, we write the system of first order differential equations governing the problem as [13],

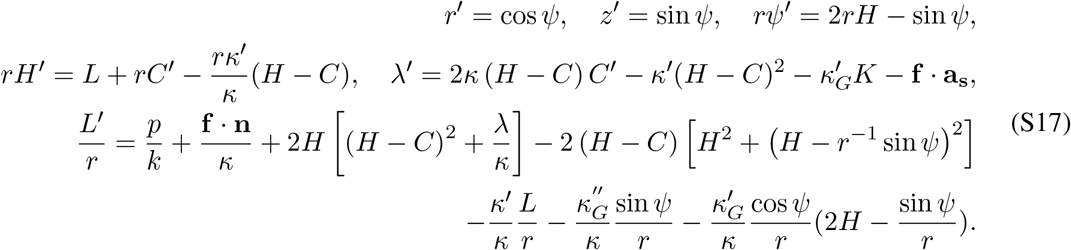

The applied boundary conditions are

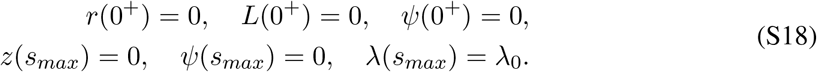

In asymmetric coordinates, the manifold area can be expressed in term of arc length [14, 15]

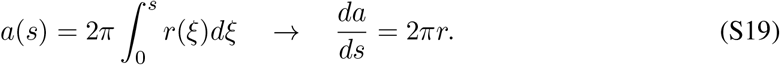

Eq. S19 allows us to convert Eq. S17 to an area derivative and prescribe the total area of the membrane.

We non-dimensionalized the system of equations as

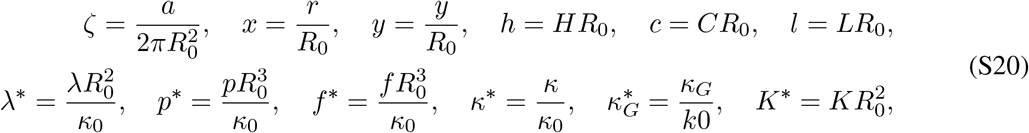

where *R*_0_ is the radius of the initially circular membrane patch.

Rewriting Eq. S17 in terms of Eq. S19 and the dimensionless variables Eq. S20, we get [13]

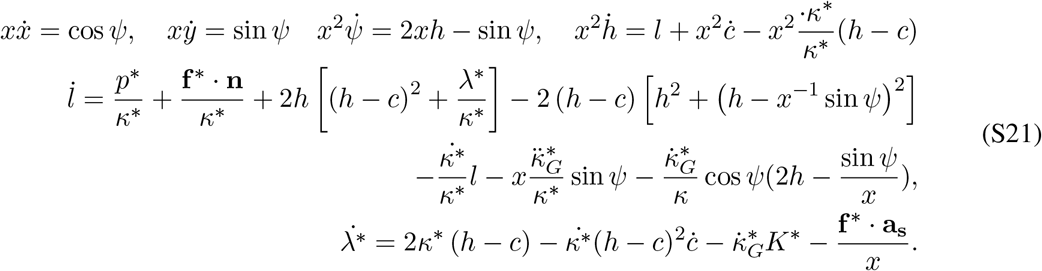

The spontaneous curvature field is modeled by a hyperbolic tangent functional as

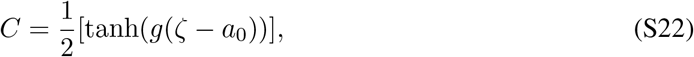

where *a*_0_ is the area of applied spontaneous curvature and *g* = 20 is a constant that ensures a sharp but smooth transition.

#### 2.3.2 Force balance along the membrane for isotropic spontaneous curvature

##### (i) Constant bending and Gaussian moduli

A general force balance for a surface *ω*, bounded by a curve *∂ω*, is (Fig. 2)

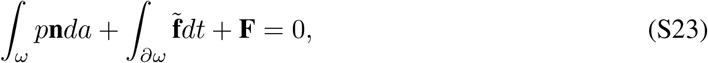

where *t* = *r*(*s*)*θ* is the length along the curve of revolution perimeter, p is the pressure difference across the membrane, 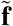 is the traction along the curve of revolution t and **F** is any externally applied force on the membrane. Along any circumferential curve on the membrane at constant z, the traction is given by [6, 11]

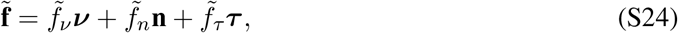

where

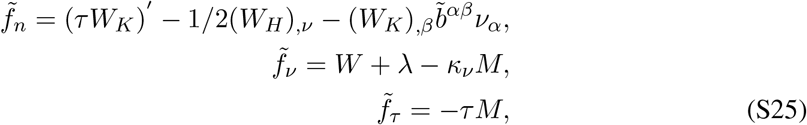

and 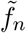, 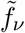 and 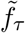 are force per unit length acting along the normal **n**, tangent *ν* to the surface, and transverse tangent **e**_*θ*_ respectively. In Eq. S25, *M* is the bending couple given by

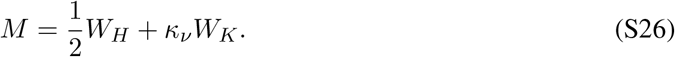

Because *τ* = 0 (no twist) in asymmetric coordinates, the normal and tangential tractions become

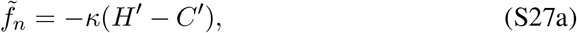

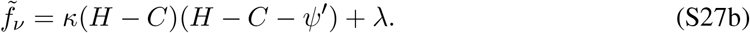

Projecting Eq. S24 onto the orthogonal bases **e**_*r*_ and **k** gives us the equation for axial and radial tractions [6, 11],

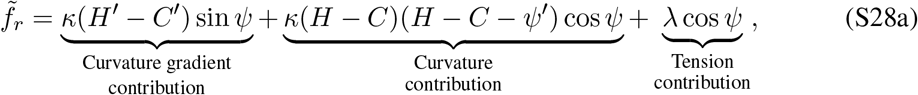

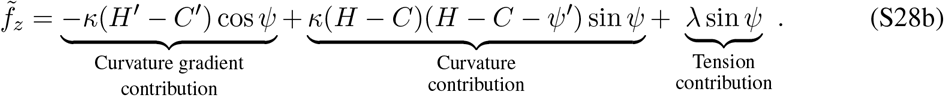

Because 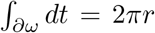, the applied force in the axial direction can be evaluated by substituting Eqs. S28a, and S28b into Eq. S23,

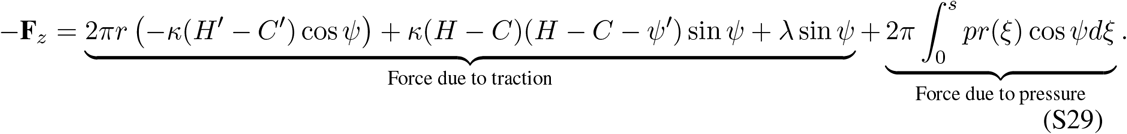

This can be rewritten in terms of tractions as

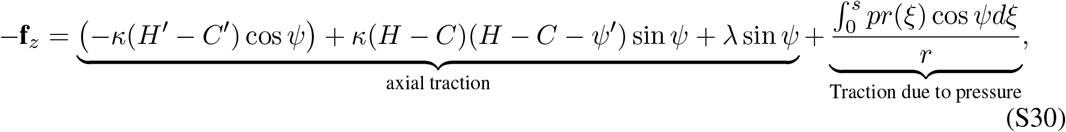

where 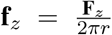. The energy per unit length *ξ*, or “effective line tension,” can be found by integrating Eq. S28a along the perimeter boundary *∂ω*,

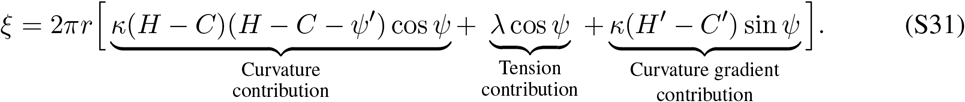

##### (ii) Spatially heterogenous bending and Gaussian moduli

For a membrane with a spatially heterogenous bending and Gaussian moduli, the normal and tangential tractions in Eqs. S27a, S27b become

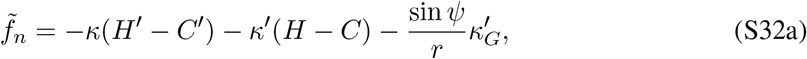

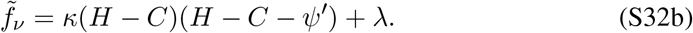

The radial and axial tractions in Eqs. S28a and S28b can be rewritten for the general case as

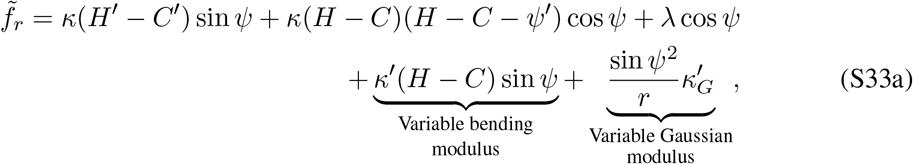

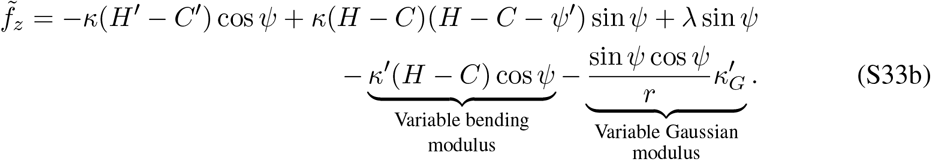

Similarly, the axial force and energy per unit lengths in Eqs. S29, S31 can be rewritten as

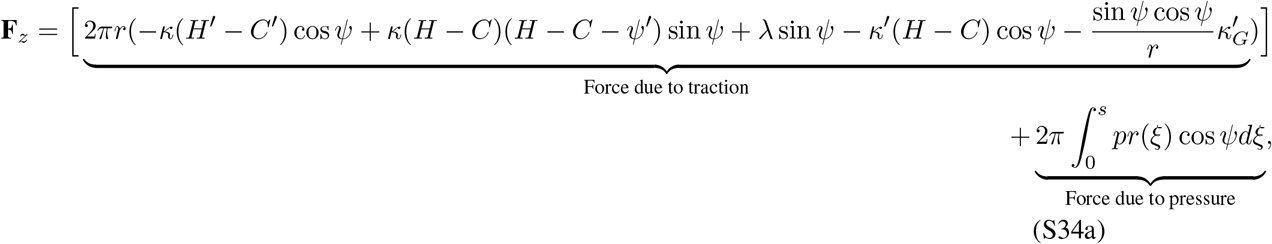

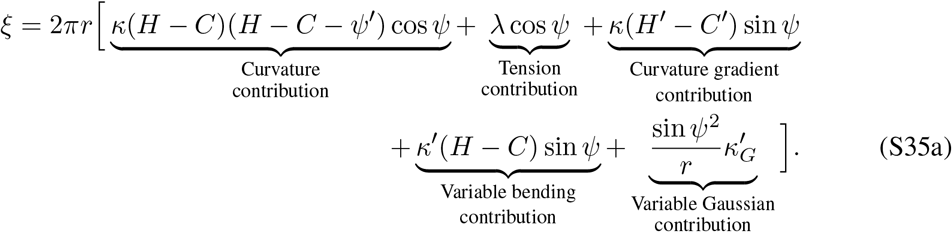

#### 2.3.3 Equation of motion for anisotropic spontaneous curvatures

By using the surface parametrization Eq. S14, we are able to define the curvature deviator (D) as

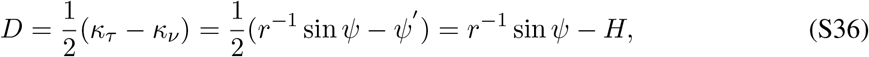

Here, we need to revise our defined L as 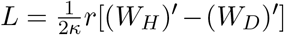, therefore for uniform bending and Gaussain modulii, the system of first order differential equations modify as [8],

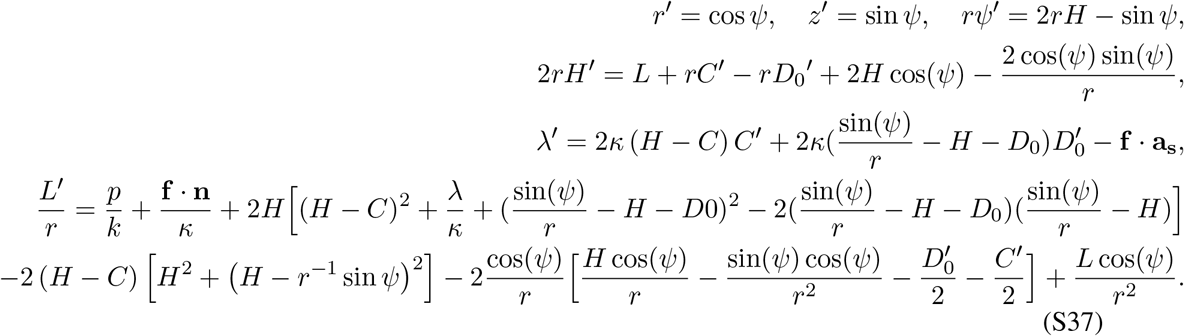

#### 2.3.4 Force balance along the membrane for anisotropic spontaneous curvatures

By considering the anisotropic spontaneous curvature contribution to the strain energy S11, the traction components in Eq. S25 and bending couple in Eq. S26 are modified

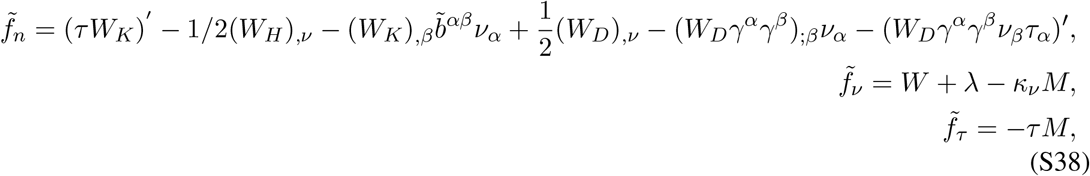

and

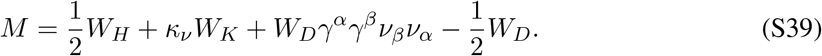

In asymmetric coordinates, the normal and tangential tractions simplify as

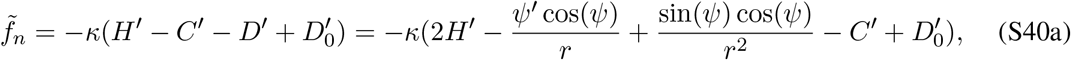

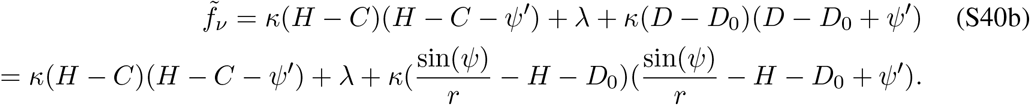

Using Eqs. S38 to simplify the traction equations

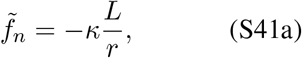

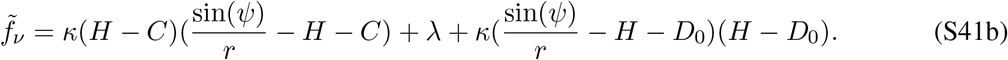

Axial force can then be written as

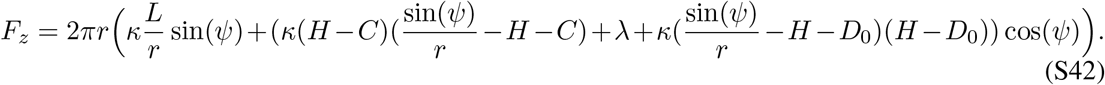

### 2.4 Asymptotic approximation for small radius

To ensure continuity at the poles, we use *L* = *H*’ = 0 as a boundary condition in our simulations. However, this boundary condition reduces the expressions for tractions (Eqs. S28b, S28a) to zero at the pole. To avoid this discrepancy, we derive an asymptotic expression for tractions at small arc length. We proceed by assuming that the pole in Eq. S21 is at *x* = 0 and choose a rescaled variable given by

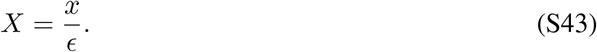

Here, *ϵ* is a small parameter, so that *X* is order of one. We can extend this to other small variables in Eq. S21 near the pole to get

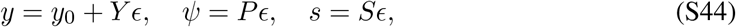

where *Y, P, S* are the corresponding rescaled parameters and *y*_0_ is membrane height at the pole.

In the simple case with no spontaneous curvature (*C* = 0), no external force **f** =0 and no pressure difference *p* = 0, we substitute Eqs. S44 and S43 into Eq. S21 and use a Taylor expansion to get

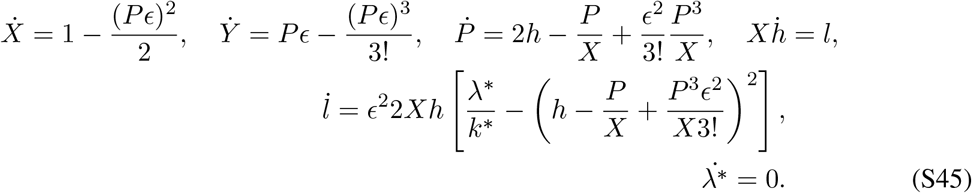

We look for a solutions with form of

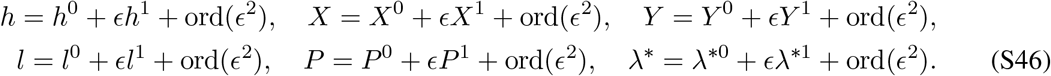

The leading order terms in Eq. S46 are

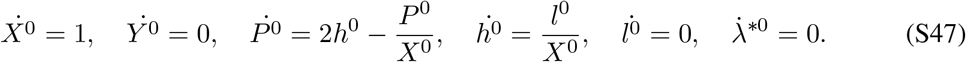

Integrating the differential equations in Eq. S47, we get

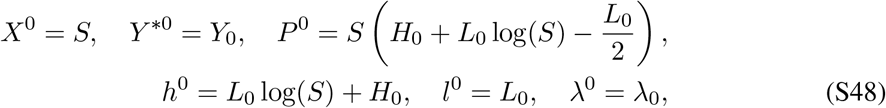

where *Y*_0_, *H*_0_ and *L*_0_, λ_0_ are integration constants. We then look at order *ϵ*^1^ terms in Eq. S45

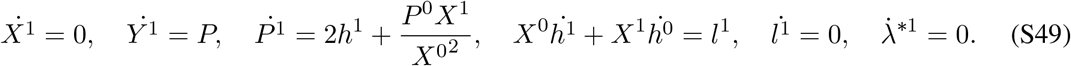

The first order terms are thus given by

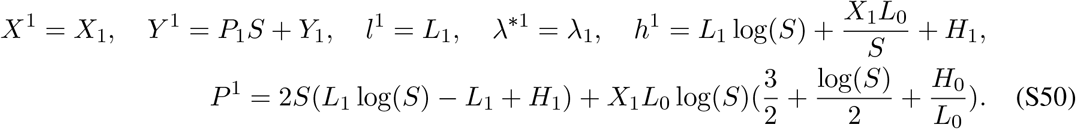

Combining the leading order and first order terms and substituting into Eq. S46, our system of variables can be written as

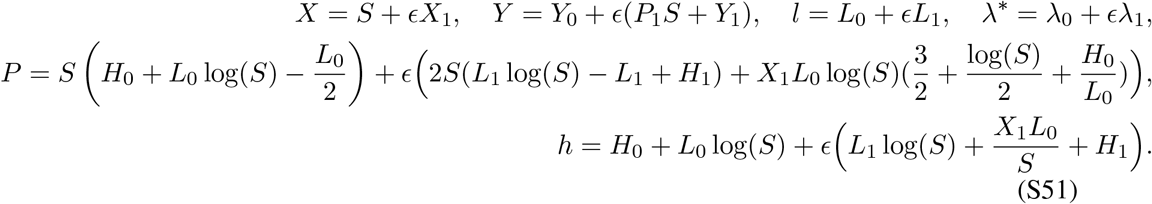

We are interested in the asymptotic expansion of mean curvature near the pole, which is given by

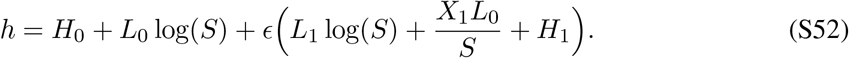

This can be rewritten as

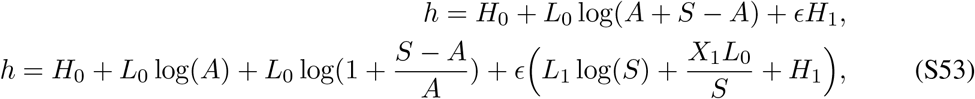

where A is a constant. If 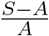 is small, we can perform a Taylor expansion around *S* = *A* to get the leading order

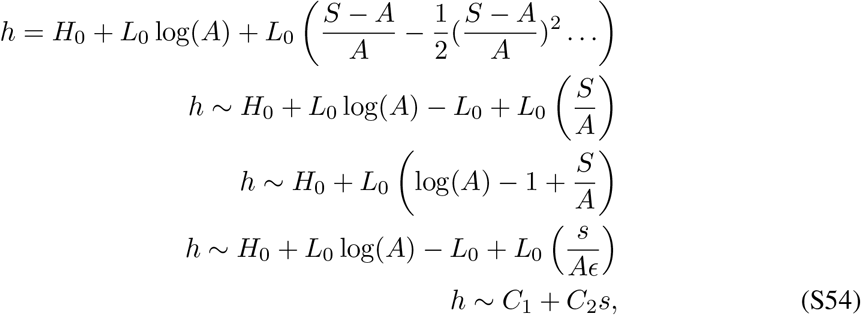

where *C*_1_ and *C*_2_ are constants. This shows that the mean curvature can be approximated as a linear solution near the pole for *S* ∼ *A* or *s* ∼ *Aϵ*. In our image analysis, inaccuracies near the pole begin at orders of magnitude of 10^−2^. At this range, we can approximate a linear solution for mean curvature. Similarly, we consider an asymptotic expansion for *ψ* near the pole at leading order

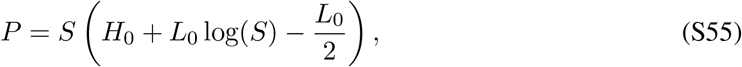

which can be rewritten as

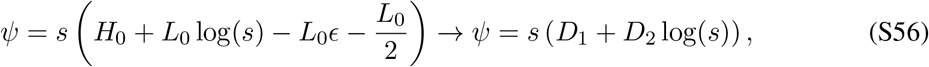

where *D*_1_ and *D*_2_ are constants. We can now substitute the approximation for mean curvature and *φ* near the pole into Eq. S28a and S28b to get

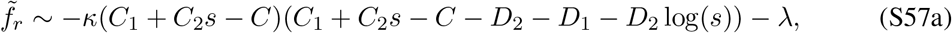

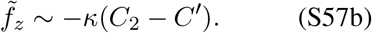

Using 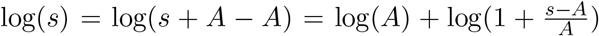 and expanding around *s* ∼ *A*, Eq. S57b can be simplified to

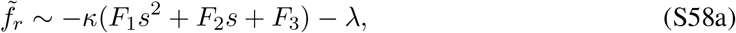

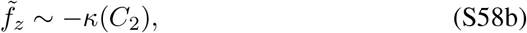

where *F*_1_, *F*_2_ are constants. We can thus approximate radial traction as quadratic in arc length near the pole, while axial traction can be correspondingly approximated as constant. In this work, we choose to start the asymptotic solution at the local minimum of mean curvature near the pole, which is *ϵ* ∼ 0.1.

## 3 Additional tether and bud formation simulations

### 3.1 Tubes pulled against pressure

In Fig. 3, we set *p* = 0 and λ_0_ = 0.02 pN/nm. However, pressure plays an important role in tether formation in certain biological contexts and thus cannot be ignored [16]. We investigated the role pressure plays during tether formation by finding pressure that produces a tube of similar radius to that obtained in Fig. 3. To do this, we first define a natural length scale for the system, *R*_0_, by the expected equilibrium radius of a membrane tube obtained by minimization of the free energy of the membrane [17].

In absence of the pressure, external force, spontaneous curvature and Gaussian modulus, we can write the free energy of the membrane as

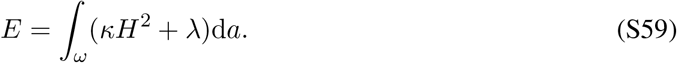

For a tube of length *L* and radius *R*, the free energy, ignoring the mean curvature of the cap 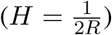, can be written as

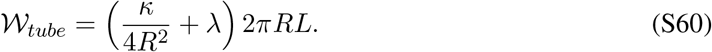

The balance between the surface tension, which acts to reduce the radius, and the bending rigidity sets the equilibrium radius *R*_0_. Taking 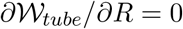 we obtain

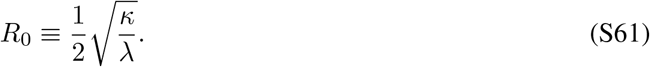

We can perform a similar analysis with pressure replacing surface tension. The free energy of the membrane Eq. S59 can be rewritten as

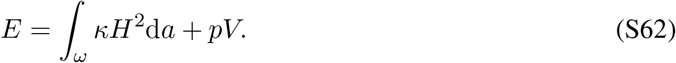

Again for a tube of length *L* and radius *R*, the free energy can be written as

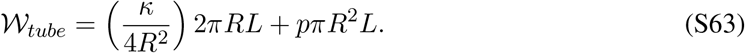

Here, the balance between pressure, which acts to reduce the radius, and the bending rigidity sets the equilibrium radius *R*_0_. Taking 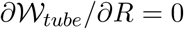 we obtain

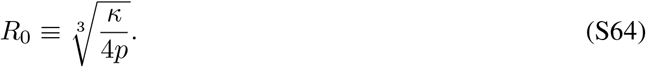

Comparing Eq. S64 and Eq. S61, we can find an equivalent pressure to the surface tension needed for achieving a tube of radius *R*_0_,

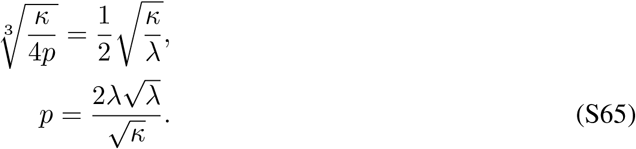

Eq. S65 gives an equivalent pressure *p* = 0.3kPa for a surface tension of 0.02pN/nm. We perform the tether pulling simulation for this value of pressure, such that the pressure acts inward for every non-zero height. Surface tension is set to zero at the base. Using the expressions for traction incorporating pressure (Eqs. S4, S5, S28), we can plot the axial, radial (Fig. S2A and B), normal and tangential tractions (Fig. S2D and E). The traction distributions show similar behaviour to Fig. 3 in the main text. Using Eq. S29, the applied force matches the difference between pressure force in the axial direction and the force due to the axial traction (Fig. S2F). Panel C shows that energy per unit length has a similar behavior to the trend in Fig. 3D of the main text.

### 3.2 Tubes pulled against pressure and surface tension

Yeast endocytic buds experience a very large pressure on the order of 1MPa [16, 18]. This large hydrostatic pressure intrinsically represents the effect of the matrix during endocytosis. In Fig. S3, we perform a tether pulling simulation for pressure 1MPa, surface tension 0.02 pN/nm and bending modulus of 32000 pN · nm, suggested by [16]. Fig. S3A and B show the axial and radial tractions for four membrane shapes as the tether is pulled out. Because of the large pressure, the radius of the tether is very small. A consequence of the small radius is a positive radial traction at the neck, where the membrane wants to push out. Axial and radial traction are both constant over cylindrical parts of the tether. Energy per unit length (seen in Fig S3C) shows large negative values at the neck and near the pole, similar to cases before. The normal and tangential traction distributions (Fig. S3D and E) along the membrane and are also qualitatively similar to the previous cases, but differ in magnitude due to larger bending modulus, with tractions being almost two orders of magnitude larger. In Fig. S3F, the external force is plotted vs the height of the tether and matches the difference between pressure force and axial force (Eq. S34a). In the presence of pressure, a much larger force is required to pull out the tube – the maximum force is almost 600 times larger than the case without pressure.

### 3.3 Axial and radial tractions in bud formation

The axial and radial tractions for the budding simulation, Fig. 5 of the main text, are shown in Fig S4. The axial traction along the membrane is negligible in all the stages of bud formation (Fig. S4A). The axial force due to traction Eq. S29 depends on three different terms, curvature, curvature gradient and surface tension. The calculated axial force at the interface is zero because tension term cancels out the force due to curvature gradient and the force associated with curvature is zero by itself (Fig. S4B). This means that neck formation is purely regulated by radial stresses (Fig. S4C). For small deformations, the radial traction is positive throughout, which shows that the membrane works to oppose the deformation. However, with the formation of U-shaped caps, radial traction changes sign and acts inward, representing the membrane’s tendency to form small necks.

### 3.4 Bud formation with a rigid protein coat

In Fig.5, we assumed the bending rigidity is homogeneous all along the membrane. However, various force microscopy measurements have shown that bending rigidity along the protein coat is much larger than the bare membrane [19]. To investigate the effect of spatial heterogeneity in the bending moduli on bud morphology and traction distributions, we repeated the simulation in Fig. 5, assuming that the bending rigidity along the spontaneous curvature field is 7.5 times larger than the bare membrane (*κ*_coat_ = 7.5 × *κ* = 2400 pN/nm (Fig. S5A)). Comparing these shapes to those in main text (Fig. 5), we see that in this case, a larger spontaneous curvature is required to form a narrow neck because the membrane is stiffer and harder to bend. Using Eqs. S32a, S32b and S35a, the normal, tangential tractions and energy per unit length distributions are plotted along the shapes (Fig. S5B, C and D). The positive normal traction distribution indicates the resistance of the membrane to bending. Similar to the case of a homogeneous membrane, the line tension can be divided into two sections. In the first section (small spontaneous curvature), tent shaped buds form and the line tension changes sign from positive to negative. Here, the average difference in the line tension magnitude between the two cases is less than 1 pN. However, in the second section (large spontaneous curvature), the line tension magnitude decreases with the formation of an Ω-shaped bud, and the average difference in the line tension magnitude is signifcantly larger (∼ 4 pN) (Fig. S5E).

### 3.5 Bud and tube formation in arc length

In Figs.5 and 3, we fixed the total area of the membrane and increased the magnitude of the spontaneous curvature and applied force respectively (Eqs. S21). However, in active non-equilibrium processes such as endocytosis, the available membrane area can vary. One possible way to consider the impact of the membrane area adjustment is solving the equations in arc length (Eqs. S17) instead of area. Here, we repeated the simulations of Figs. 5 and 3 for a fixed arc length of membrane. In the case of the bud, we fixed the arc length coverage of the coat and increased the spontaneous curvature from 0 to C=-0.032 nm^−1^ (Fig. S6A). The tractions and energy per unit length distribution along different shapes are shown in Fig. S6B-D. Evidently, independent of whether the available area of the membrane is fixed or not, the energy per unit length at the interface is between 6 to −6 pN and changes sign from positive to negative with formation of a neck (Fig. S6E). In the case of a tube simulation for a fixed arc-length (Fig. S7), we can replicate the force-displacement curve in the main text (Fig. S7B and Fig. 3B), which is for a fixed membrane area. The tractions and energy per unit length distribution along a few shapes are shown in Fig. S7A,C,D.

### 3.6 Bud formation with cytoskeleton forces

We consider the effects of the cytoskeleton during endocytosis as previously described in [13]. In Fig. S8, we apply actin-mediated forces on a U-shaped bud such that actin polymerizes in the form of a ring at the base of the endocytic pit with the network attached to the protein coat [13, 20]. Fig. S8B-D show the tractions and energy per unit length distributions along the initial and final shapes as the membrane is pulled out. Fig. S8E shows the match obtained between the applied force and axial force calculated at the edge of the protein coat. Here, we note that calculating the axial force at the base predicts zero applied force – a consequence of the actin ring acting at the base that integrates to zero. To further emphasize this point, we repeated the simulation without this downward force at the base (Fig. S9) and show that the match between applied force and axial force can be obtained both at the base of the membrane and at the edge of the protein coat. Fig S9B shows that the large tangential traction along the neck is limited to a smaller region (red) compared to Fig. S8B, because without considering the actin ring force at the base, there is lesser axial stretch along the bud neck.

### 3.7 Bud formation with anisotropic spontaneous curvature

Proteins induce a highly anisotropic local spontaneous curvature [9, 10]. To model this effect, we used a modified energy functional (Eq. S11) that includes deviatoric curvature effects. This then can be written as the shape equation (Eq. S12) and tangential variation equation (Eq. S13). We solve this system of equations for a deviatoric curvature field applied over the cylindrical portion of a membrane tube (Fig. S10, [20]). Fig S10A shows neck formation in a membrane tube with increasing deviatoric curvature. The membrane invagination obtained resembles the PM shape seen during assembly of rvs proteins at the neck of a tube [20]. We note here that we apply both a spontaneous curvature and deviatoric curvature with opposite signs. Fig. S10D shows that the energy per unit length at both interfaces matches the trend seen in Fig. 6 leading up to neck formation. Fig. S10E shows that the axial forces can be matched to applied forces. Axial forces decrease as a consequence of the membrane height being constrained.

### 3.8 Additional sensitivity analysis

Fig. 8B shows that the axial force (F_*z*_) is not sensitive to error in membrane tension for a fixed shape of a membrane tether. This is because the calculation of axial force was performed at the base of the PM invagination where the membrane is nearly flat (*ψ* = 0). In Fig. S11, we perform the same analysis by repeating the calculation at two other locations – (1) at the edge of the area of applied force, (2) at the point of zero mean curvature. Fig. S11A shows these points (1) and (2) along the membrane shape corresponding to the peak of the force-displacement graph in Fig. 3B. The error in (F_z_) due to membrane tension increases for calculations at regions of larger tangent angle *ψ*(Fig. S11B, C). Thus, to minimize error in axial force, we choose to perform traction calculations at base of the membrane.

## 4 Supplementary figures

**Figure S1:**
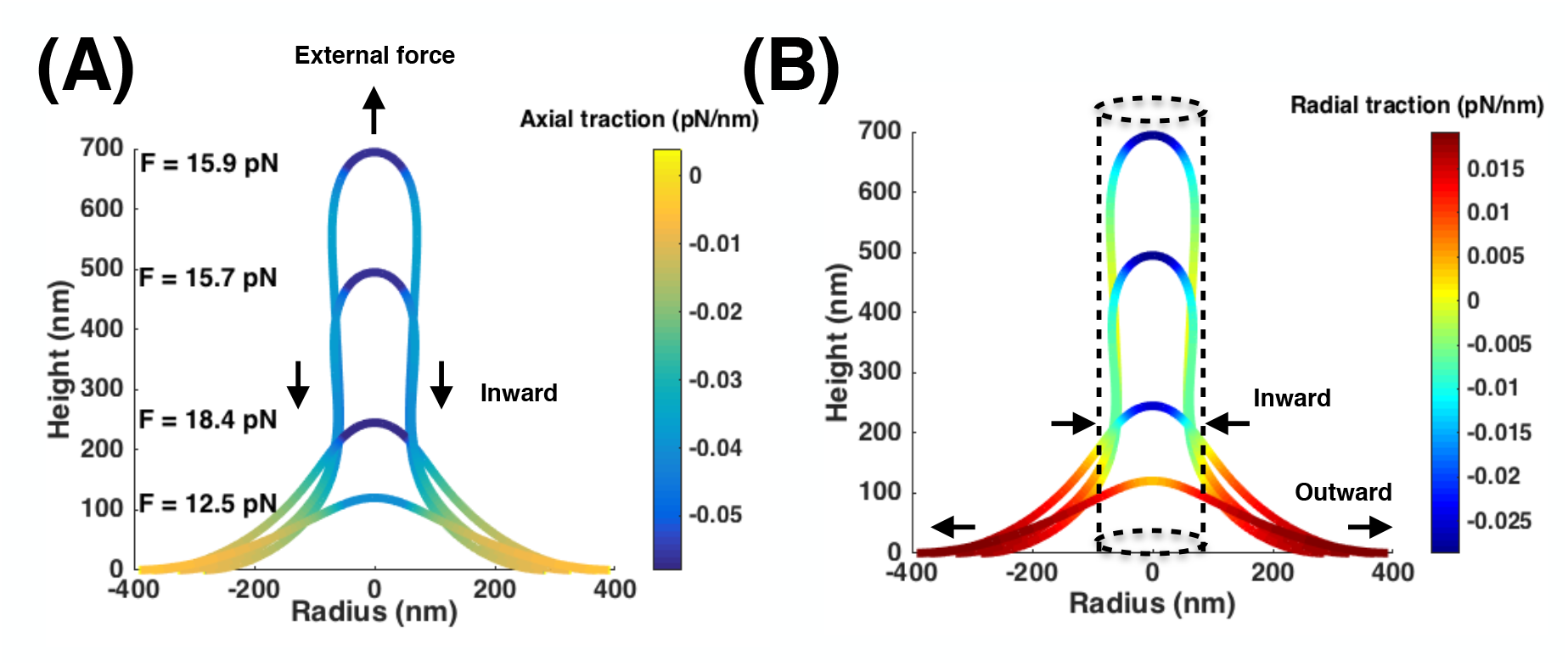
Axial and radial traction (Eqs. S28b, S28a) distribution plotted along the same membrane shapes as in Fig. 3A–C. (A) Axial traction distribution. The axial traction is constant along the tube. (B) Radial traction distribution. The dotted line is the stable cylindrical geometry.

**Figure S2:**
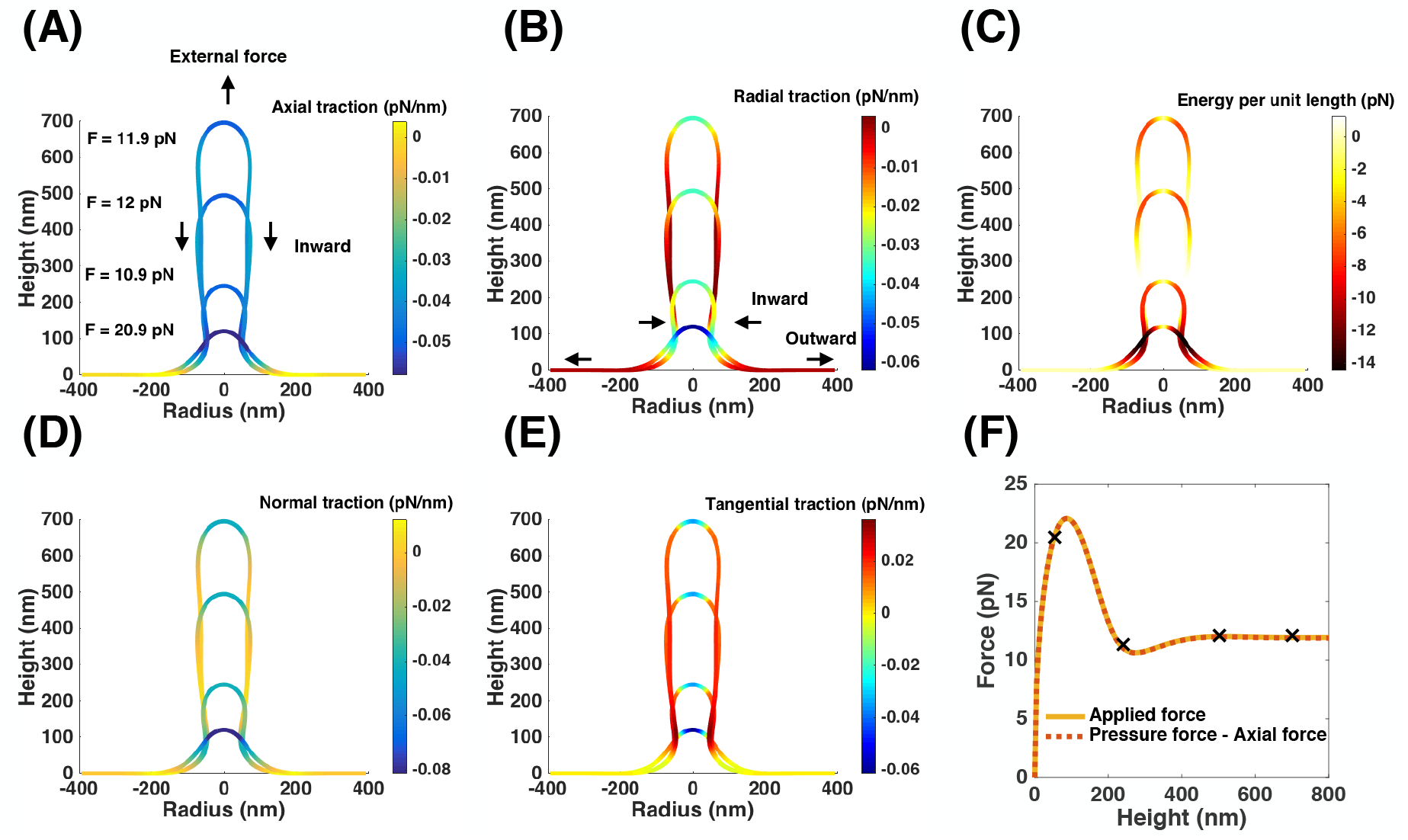
Tether pulling simulation for the pressure of 0.3kPa, bending modulus 320 pN · nm, no surface tension at the boundary (λ_0_ = 0), and a point force. (A) Axial traction distribution along the tether. (B) Radial traction distribution. We find a negative value at the neck and a positive value at the base. (C) Energy per unit length (Eq. 6) plotted along the shapes. We observe a large value at the neck - predicting an ‘effective’ line tension of 11 pN for a tether of height 700 nm. (D) Normal traction distribution. It is large and negative over the area of applied force. (E) Tangential traction distribution. (F) Applied force and the difference between the calculated pressure and axial force (Eq. S29) plotted as a function of tether height.

**Figure S3:**
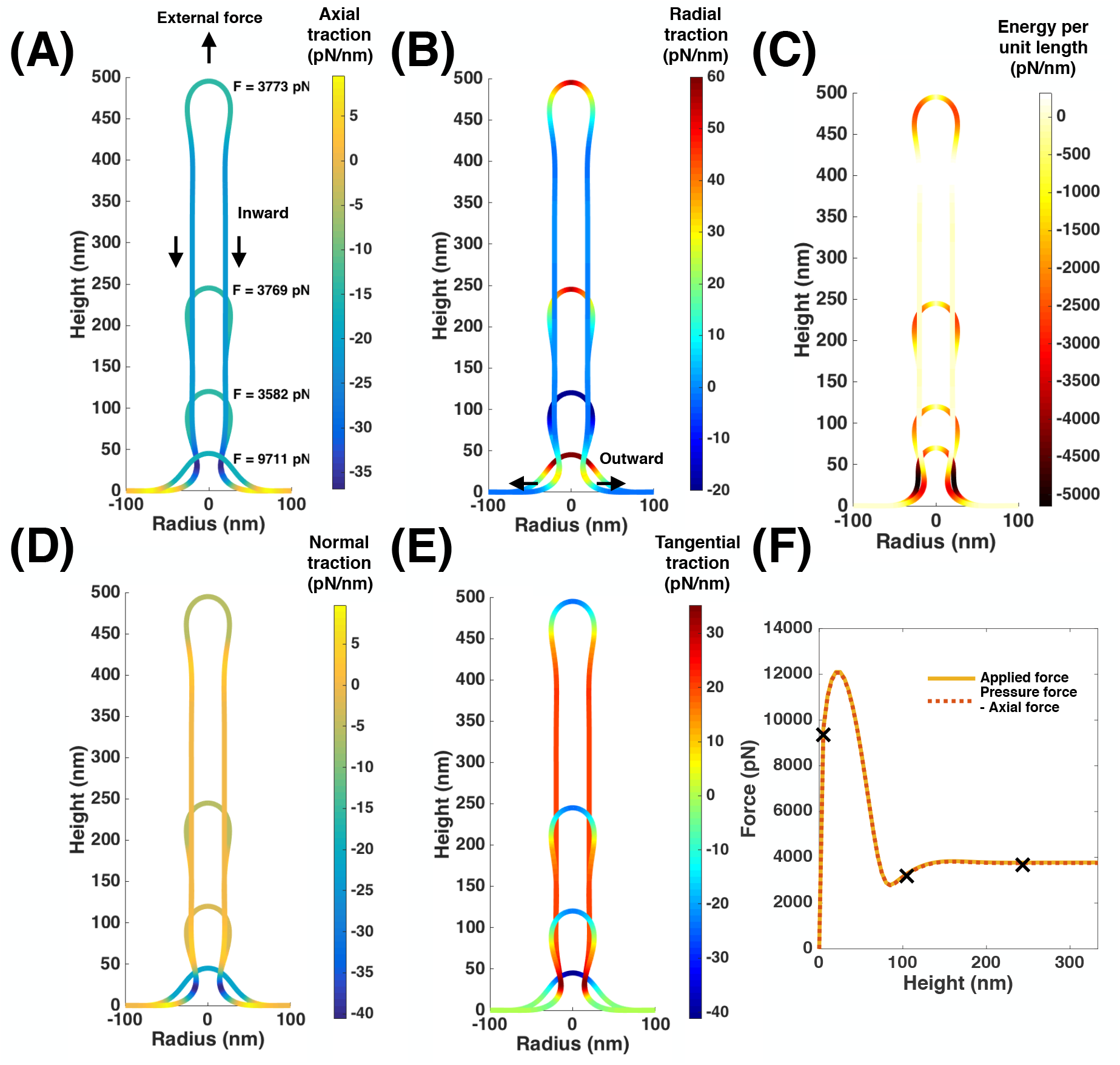
Tether pulling simulation for a pressure of 1MPa, bending modulus 32000 pN · nm, surface tension at the boundary (λ_0_ = 0.02pN/nm), and a point force. (A) Axial traction distribution along the tether for four chosen membrane shapes. (B) Radial traction distribution along the membrane shapes in (A). (C) Energy per unit length (Eq. S31) plotted along the membrane shapes in (A). We observe a large value at the neck. (D) Normal traction distribution along the membrane shapes in (A). It is large and negative over the area of applied force and at the neck. (E) Tangential traction distribution along the membrane shapes in (A). (F) Applied force and the difference between the calculated pressure and axial force (Eq. S29) plotted as a function of tether height.

**Figure S4:**
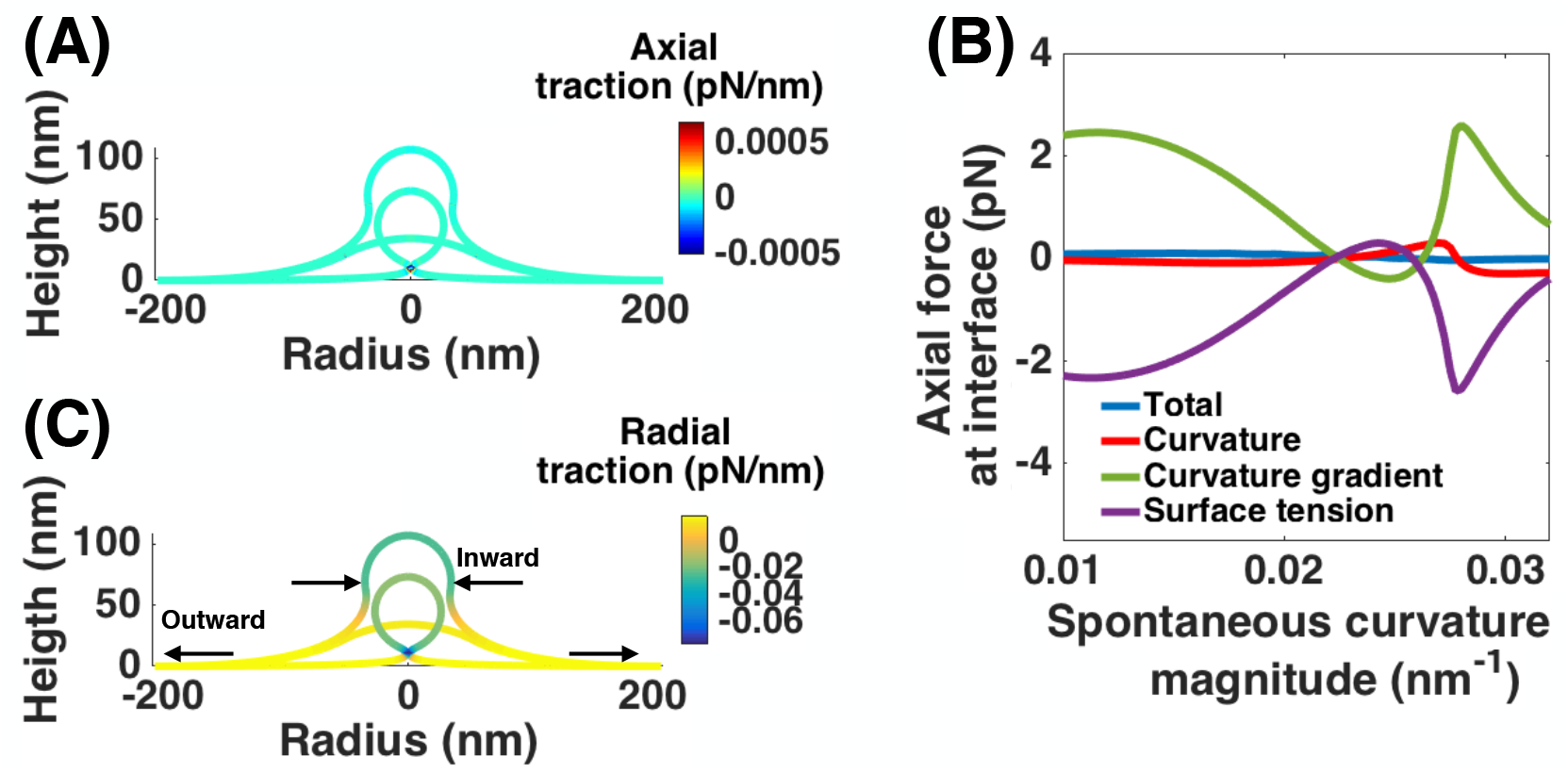
Bud formation from a flat membrane for increasing spontaneous curvature and a constant area of spontaneous curvature field A = 10,053 nm^2^. The spontaneous curvature magnitude increases from *C* = 0 to *C* = –0.034 nm^−1^, the bending modulus is *κ* = 320 pN · nm and surface tension at the edge is λ_0_ = 0.02 pN/nm. Axial traction does not play any role in invagination. (A) Axial traction along the membrane is negligible for all shapes. (B) Axial force at the interface is almost zero. Terms due to tension and curvature gradient cancel each other and force due to curvature is automatically zero. (C) Radial traction distribution for three shapes. A large negative radial traction at the neck should favor membrane scission.

**Figure S5:**
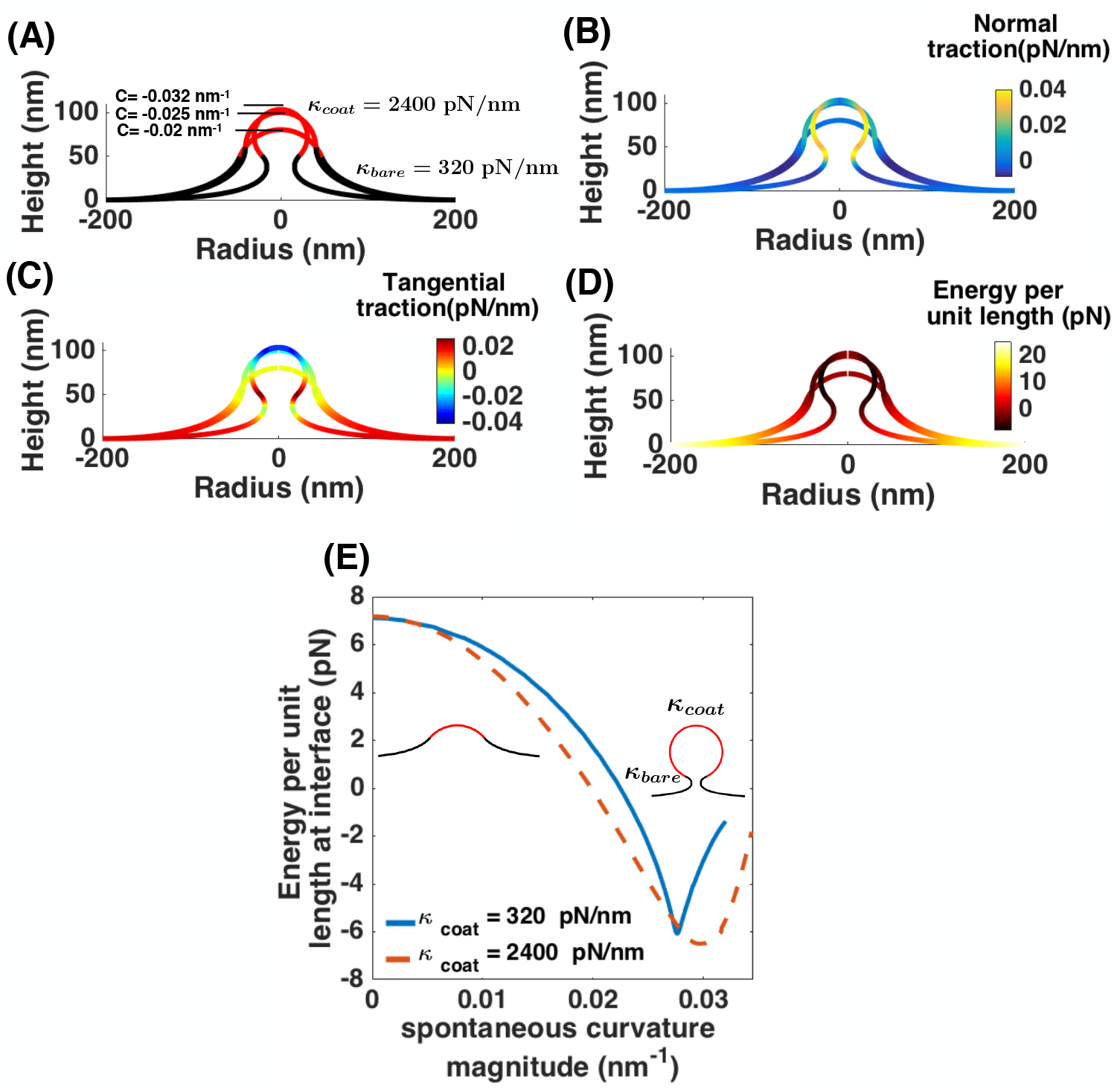
Analysis of membrane budding due to protein-induced spontaneous curvature with a rigid coat. Simulations for Fig. 5 is repeated for *κ*_coat_ = 2400 pN/nm and *κ*= 320 pN/nm. (A) Membrane shapes for same three spontaneous curvature as Fig. 5 A. (B) Normal traction along the membrane for the shapes shown in (A). (C) Tangential traction distribution (D) Energy per unit length distribution for the chosen shapes. (E) Energy per unit length at the edge of the edge of the spontaneous curvature field as a function of spontaneous curvature for the homogeneous membrane in Fig. 5 (blue solid line) and a rigid protein coat (red dashed line). In large values of spontaneous curvature (Ω- shaped bud) the average difference between the line tension magnitudes is almost 4 pN.

**Figure S6:**
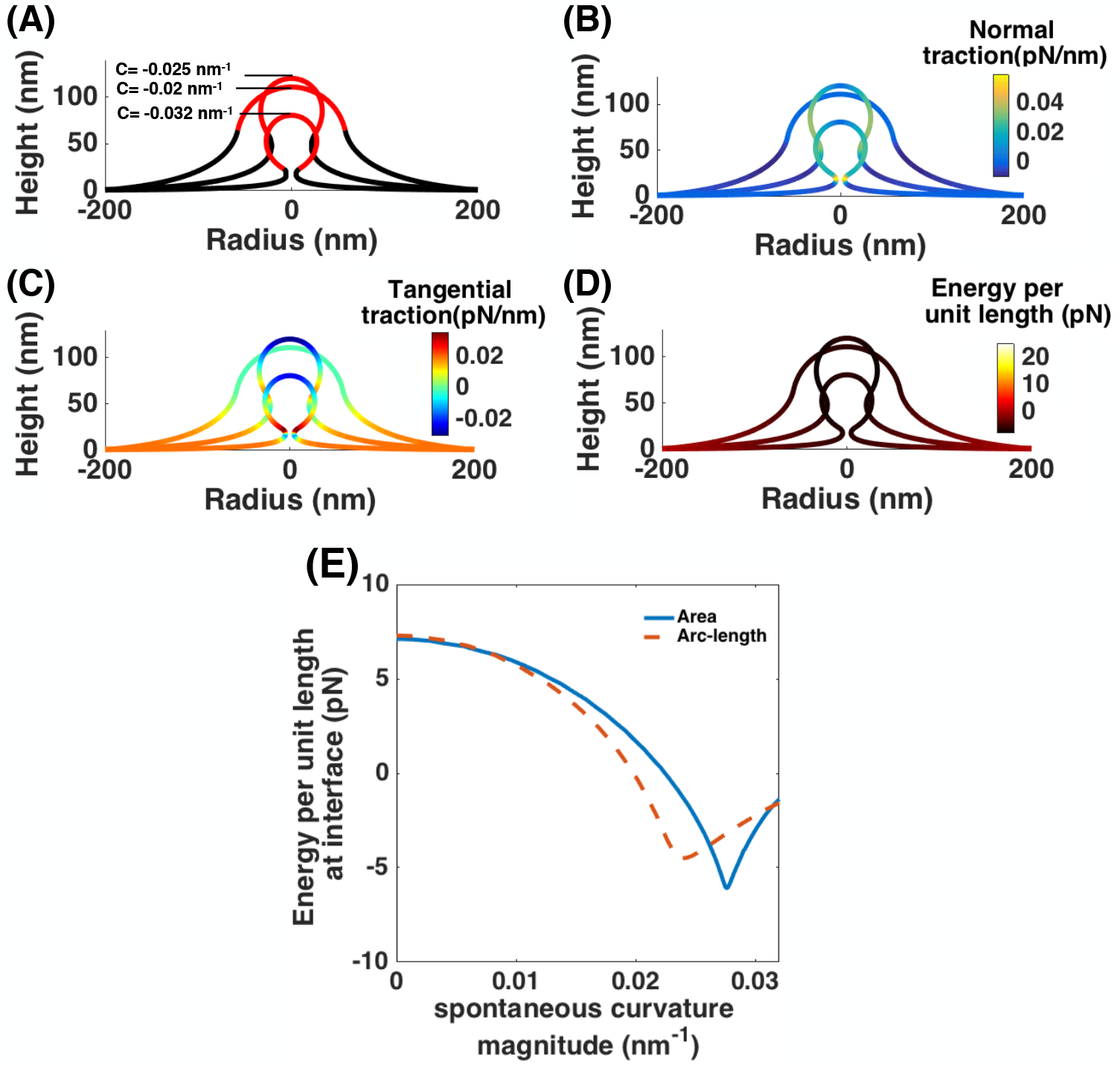
Budding simulation with protein-induced spontaneous curvature for a fixed arc length instead of a fixed membrane area. (A) Three different membrane shapes with increasing spontaneous curvature. (B-D) Normal traction, tangential traction and energy per unit length distribution along the observed shapes in panel (A). (E) Energy per unit length at the edge of the protein coat. Blue solid line is for the fixed membrane area (Fig. 6) and the red dashed line represents the case with constant arc length.

**Figure S7:**
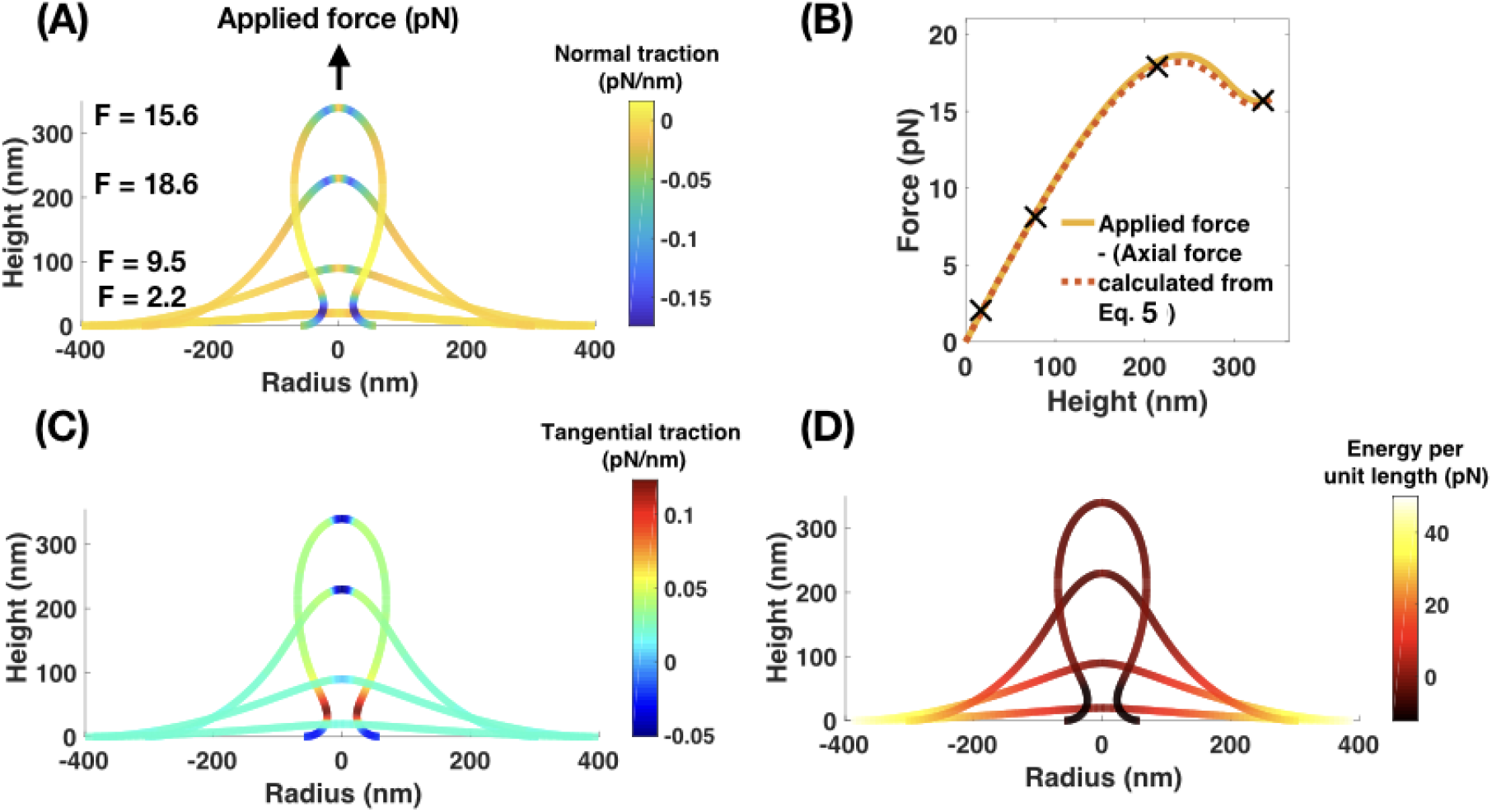
Tether pulling simulation with a point force for a fixed arc length instead of a fixed membrane area. Bending modulus *κ* = 320 pN · nm, surface tension at the boundary (λ_0_ = 0.02pN/nm). (A) Normal traction distribution along four chosen membrane shapes. (B) Match obtained between applied force and axial force calculated from Eq. S29 plotted vs height of the tether. (C) Tangential traction distribution along the membrane shapes in (A). (D) Energy per unit length (Eq. S31) along the membrane shapes in (A).

**Figure S8:**
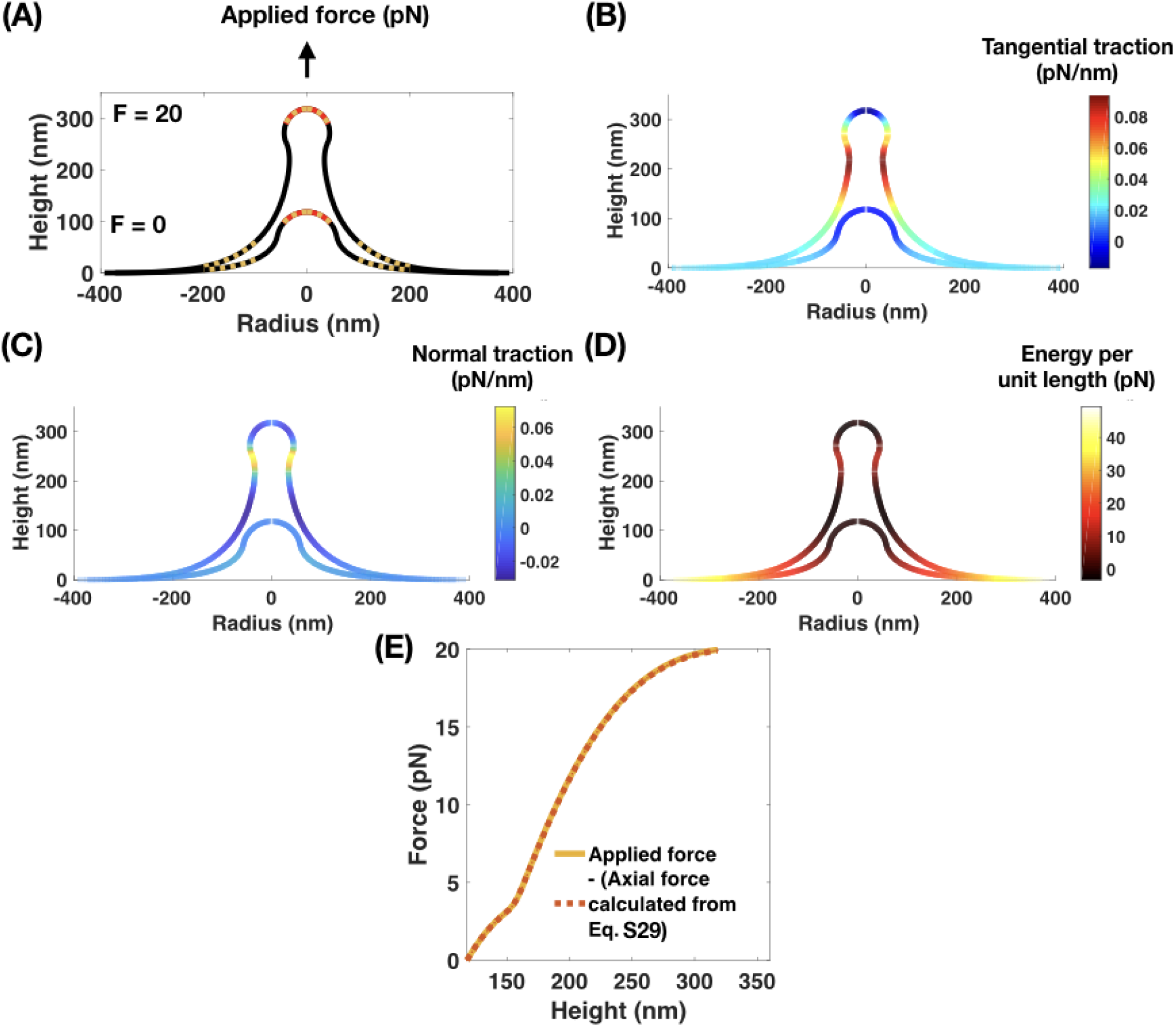
Application of axial forces (brown) onto a U-shaped bud covered by a protein coat (red). Spontaneous curvature magnitude *C* = –0.02 nm^−1^, area of spontaneous curvature field A = 17, 593 nm^2^, bending modulus *κ* = 320 pN · nm and surface tension at the edge λ_0_ = 0.02 pN/nm. Here, axial forces are applied such that there is an upward force over the protein coat and a downward force acting as a ring at the base [13]. (A) Initial and final membrane shapes obtained. (B) Tangential traction distribution along the membrane shapes in (A). (C) Normal traction distribution along the membrane shapes in (A). (D) Energy per unit length along the membrane shapes in (A). (E) Force match obtained between applied force and negative of axial force calculated using Eq. S29. Here, axial force is calculated at the edge of the protein coat. Axial force at the base is zero since the upward and downward forces balance each other out.

**Figure S9:**
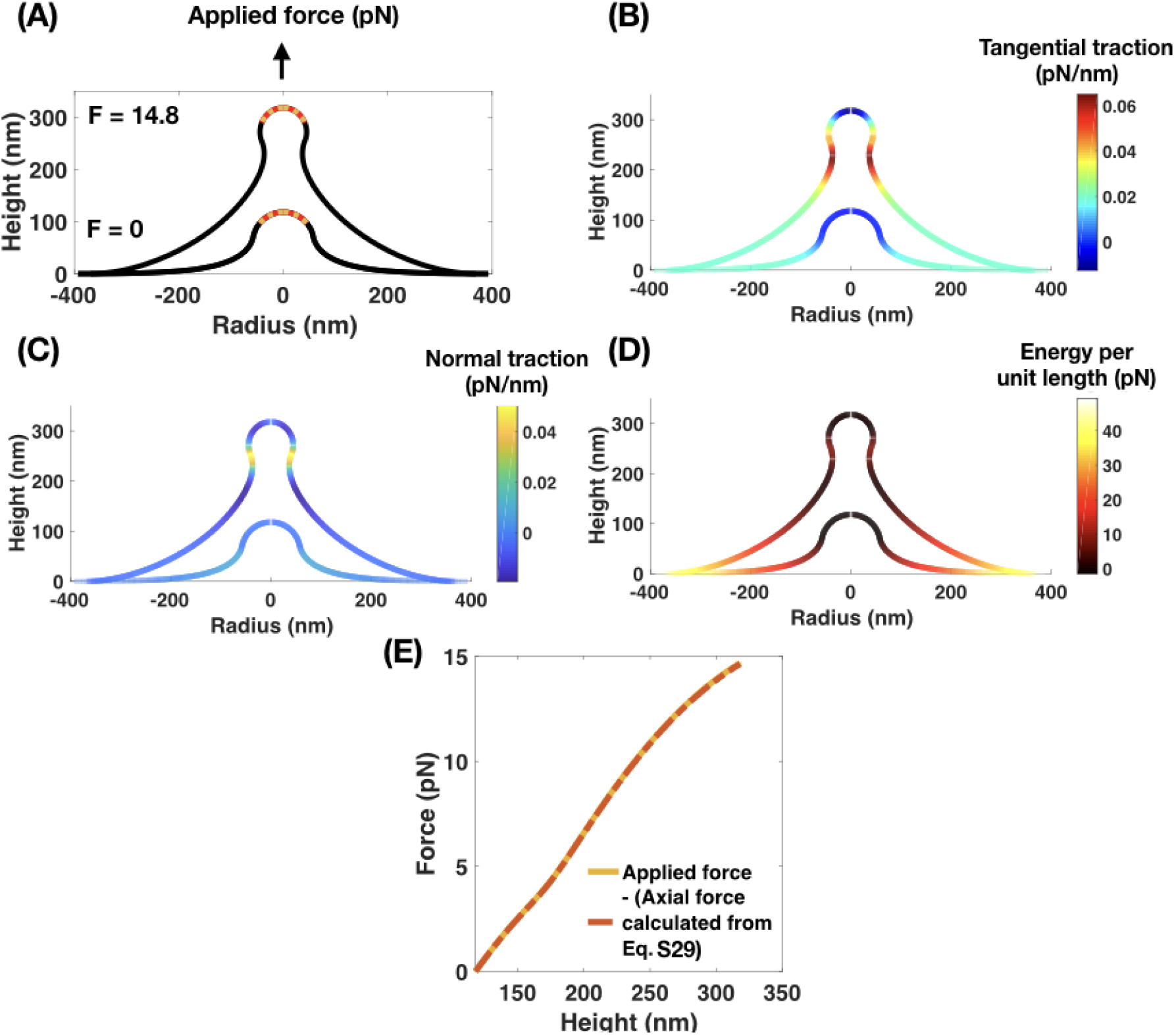
Application of axial forces (brown) onto a U-shaped bud covered by a protein coat (red). Spontaneous curvature magnitude *C* = –0.02 nm^−1^, area of spontaneous curvature field A = 17, 593 nm^2^, bending modulus *κ* = 320 pN · nm and surface tension at the edge λ_0_ = 0.02 pN/nm. Here, axial forces are applied such that there is only an upward force over the protein coat. (A) Initial and final membrane shapes obtained. Force required is smaller than Fig. S8. (B) Tangential traction distribution along the membrane shapes in (A). (C) Normal traction distribution along the membrane shapes in (A). (D) Energy per unit length along the membrane shapes in (A). (E) Force match obtained between applied force and negative of axial force calculated using Eq. S29. Here, axial force is calculate at the base of the membrane. The same match can be obtained at the edge of the protein coat.

**Figure S10:**
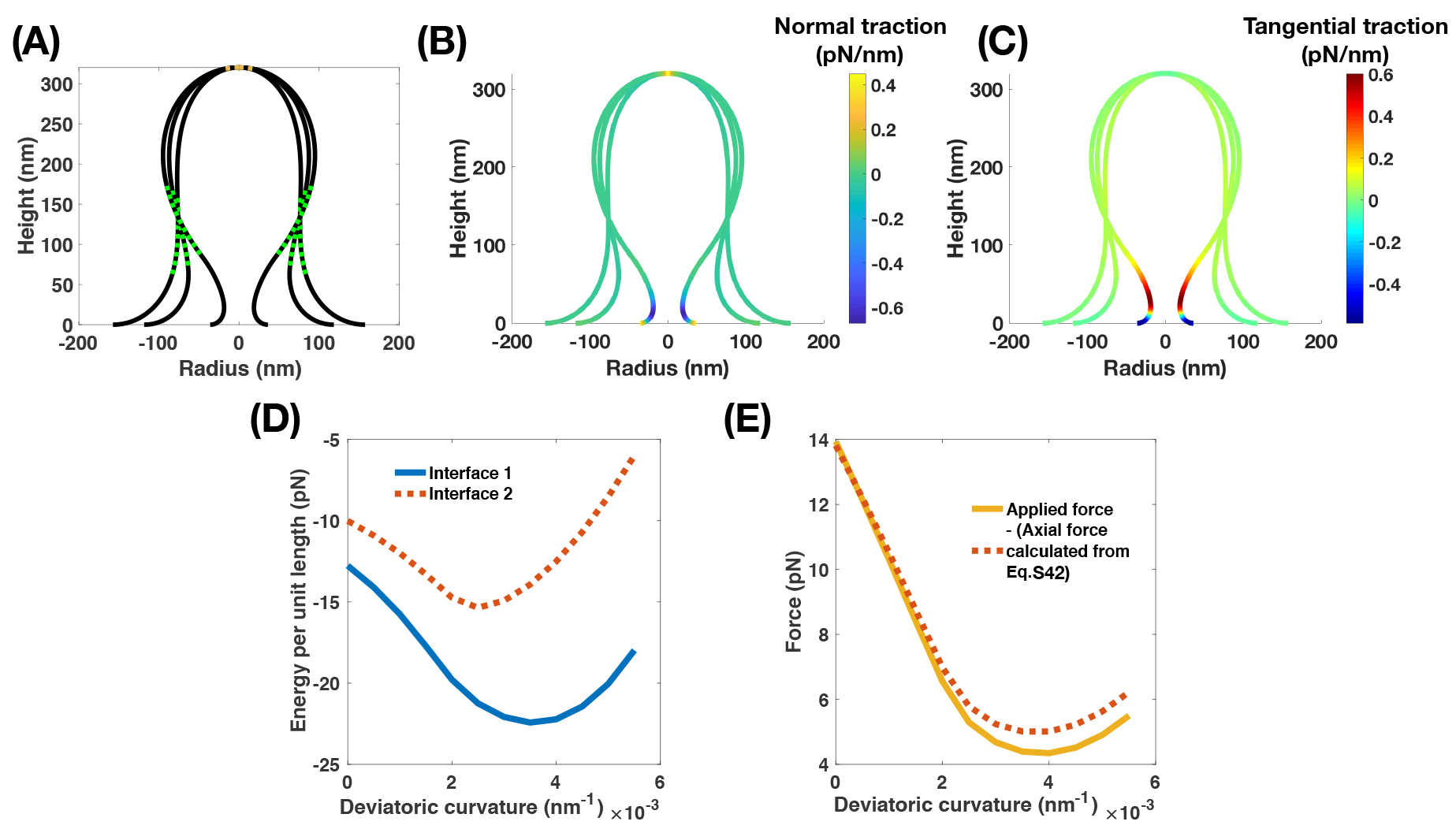
Application of a deviatoric spontaneous curvature along the cylindrical portion of a membrane tube leads to neck formation [20]. The simulation was performed by first pulling out a membrane tube of fixed arc length, followed by application of both spontaneous curvature and deviatoric curvature for a fixed height of membrane tube. Arc length of the deviatoric spontaneous curvature field s = 5 nm, bending modulus *κ* = 320 pN · nm and surface tension at the edge λ_0_ = 0.02 pN/nm.(A) Membrane shapes at a spontaneous curvature *C* = 0nm^−1^, *C* = –0.004 nm^−1^, *C* = –0.01nm^−1^ and deviatoric spontaneous curvature *D* = 0nm^−1^, *D* = 0.004nm^−1^, *D* = 0.01 nm^−1^ respectively. (B) Normal traction distribution along the membrane shapes in (A). (C) Tangential traction distribution along the membrane shapes in (A). (D) Energy per unit length at both interfaces with increasing deviatoric curvature. The trend of energy per unit length resembles the trend leading up to neck formation in Fig 5. (E) Match between applied force and axial force calculated using Eq. S42. Axial force relaxes with membrane neck formation.

**Figure S11:**
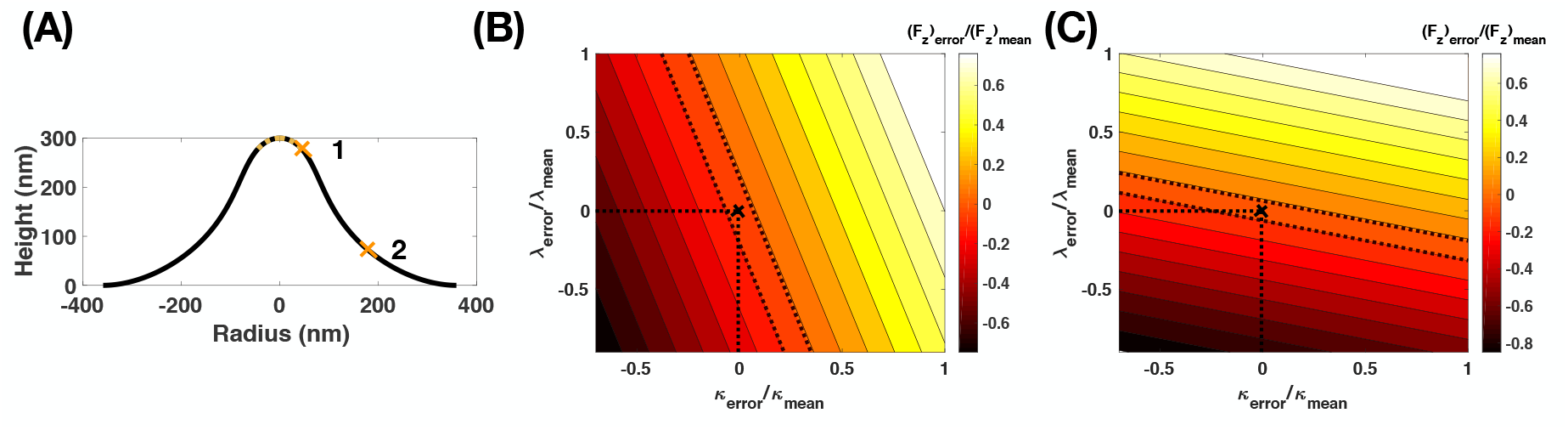
Location dependence of the sensitivity analysis to axial force calculation from a single simulation of a membrane tube. Dashed lines indicated 10 % error. (A) Membrane shape at a mean value of κ= 320 pN.nm, λ_0_ = 0.02 pN/nm, – *F_z_* (brown)= 18.0167 pN (corresponding to a tube of height 300 nm in Fig. 3). The cross marks (labeled 1 and 2) indicate the locations where axial force is calculated. (B) Error in F_*z*_ calculated at point 1 (edge of the area of applied force). (C) Error in F_*z*_ calculated at point 2 (location of zero mean curvature). Error in F_*z*_ due to error in membrane tension λ increases near the curved portions of the PM invagination.

